# Targeting posttranslational modifications of RioK1 inhibits the progression of colorectal and gastric cancers

**DOI:** 10.1101/171058

**Authors:** Xuehui Hong, He Huang, Zhijie Ding, Xing Feng, Yuekun Zhu, Huiqin Zhuo, Jingjing Hou, Wangyu Cai, Xinya Hong, Hongjiang Song, Zhiyong Zhang

## Abstract

RioK1 has recently been shown to play important roles in cancers, but its posttranslational regulation is largely unknown. Here we report that RioK1 is methylated at K411 by SETD7 methyltransferase, and that lysine-specific demethylase 1 (LSD1) reverses its methylation. The mutated RioK1 (K411R) that cannot be methylated exhibits a longer half-life than does the methylated RioK1. FBXO6 specifically interacts with K411-methylated RioK1 through its FBA domain to induce RioK1 ubiquitination. Casein kinase 2 (CK2) phosphorylates RioK1 at T410, which stabilizes RioK1 by antagonizing K411 methylation and impeding the recruitment of FBXO6 to RioK1. Functional experiments demonstrate the RioK1 methylation reduces the tumor growth and metastasis in CRC and GC. Importantly, the protein levels of CK2 and LSD1 show an inverse correlation with FBXO6 and SETD7 expression in human CRC tissues. Therefore, this study highlights the importance of a RioK1 methylation-phosphorylation switch in determining CRC and GC development.

## Introduction

Colorectal cancer (CRC) is the second leading cause of cancer-related deaths and one of the most common malignancies in the Western world (Labianca et al., 2013). Over the last decades, treatment strategies have significantly improved, however, CRC remains a high-risk gastrointestinal malignancy. And the occurrence of metastasis after operation is the leading cause of poor prognosis for CRC patients (Yoshii et al., 2014). Therefore, it is of paramount importance to decipher the potential mechanisms underlying CRC growth and metastasis, which may contribute to the development of effective therapeutics for treating CRC patients.

Rio (right open reading frame) <u>kinase</u> is a relatively conserved family of atypical <u>serine/threonine kinases</u> (LaRonde-LeBlanc and Wlodawer, 2005). RioK1 and RIOK2 are two members of the Rio family, named for yeast (*S. cerevisiae*) Rio1p and Rio2p, respectively (LaRonde-LeBlanc and Wlodawer, 2005). A third member named RIOK3 has greater similarly to RioK1, but is only known to exist within multicellular eukaryotes (Yuan et al., 2014). Although these kinases contain a kinase fold structurally homologous to eukaryotic serine-threonine protein kinase domains, they lack substrate binding loops and loops activation domain, and their actual *in vivo* targets are unknown (Mendes et al., 2015). RioK1 knockout is lethal, and was identified being an essential gene in yeast (Angermayr et al., 2002). Functionally, the yeast RioK1 orthologue affects <u>cell cycle</u> progression and <u>ribosomal</u> biogenesis (Widmann et al., 2012), and the *C. elegans* RioK1 is essential for reproduction (Weinberg et al., 2014), whereas in multicellular organisms its role remains poorly understood. Recently, several studies have reported that the RIO kinases function in RTK and PI3K signaling pathway (Read et al., 2013), and are required for the survival of Ras-dependent cancer cells (Luo et al., 2009). However, to date, no specific function has been ascribed to RioK1 in the context of human CRC and GC.

The posttranslational modification (PTM, such as phosphorylation, ubiquitination, and acetylation) of proteins is well-known to dynamically change protein function by fine-tuning protein stability, localization, or interactions (Jensen, 2006). PTMs of proteins rapidly and reversibly regulate cells in response to different stresses. Therefore, once demonstrated, these PTMs could potentially serve as therapeutic targets (Krueger and Srivastava, 2006). Among various posttranslational modifications, lysine methylation acts as a novel regulatory mechanism to control protein functions (Oudhoff et al., 2013). However, most previous studies have predominantly highlighted histone methylation, until recently accumulating evidence indicates the widespread presence of lysine methylation in nonhistone proteins (Patel et al., 2011). Although there are about 50 lysine methyltransferases in mammals, lysine methylation is primarily catalyzed by a family of protein methyltransferases containing a catalytic SET domain (Dillon et al., 2005). su(var)3-9, enhancer-of-zeste, trithorax (SET) domain-containing protein 7 (SET7), also known as SETD7, SETD9, and SETD7, that acts on histone H3K4, has been shown to monomethylate various nonhistone proteins, including Gli3, FOXO3a, p53, HIF-1α, TAF10, and Suv39h1 (Fu et al., 2006; Kim et al., 2016; Couture et al., 2006; Wang et al., 2013). This methylation produces varied results. For example, methylation of Gli3 by SET7 stabilizes Gli3, whereas methylation of HIF-1α stimulates HIF-1α proteasomal degradation (Fu et al., 2006; Kim et al., 2016). In addition, there is increasing evidence that other PTMs may parallel methylation in the regulation of various biological processes, which provide a sophisticated level of crosstalk (Bigeard et al., 2014). Although RioK1 can be regulated by phosphorylation (Angermayr et al., 2007), the enzymes responsible for other PTMs of RioK1 remain elusive.

Given that many kinases are largely regulated by PTMs (Oliveira and Sauer, 2012), we speculate that RioK1 may be regulated by methylation, or phosphorylation, or ubiquitination. The aim of this study was to investigate the PTMs and role of RioK1 in CRC and GC. Our findings definitely will further deepen the understanding of the crosstalk of PTMs signaling, expand the existing knowledge of the RioK1, and provide a novel mechanism by which RioK1 regulates tumorigenesis and metastasis.

## Results

### High-level RioK1 is closely associated with an aggressive CRC and poor overall survival

To determine the role of RioK1 in the pathogenesis of CRC, we firstly investigated the levels of RioK1 in a group of 5 CRC patient tissues and matched normal tissues. As shown in the Figure 1A, compared with the corresponding non-tumor samples, RioK1 was significantly increased in CRC tissues. The scatter diagram demonstrated the increase of RioK1 mRNA levels in CRC and metastasis lymph node samples versus normal tissues, with an average 4.03-fold and 6.15-fold increase respectively (Figure 1B). To confirm the increased RioK1 protein expression in a larger sample group, and correlate this to clinical phenotype, we performed immunohistochemical staining (IHC) on the CRC tissue array comprised of 120 patients. IHC demonstrated that CRC tissues showed higher expression of RioK1 compared to matched normal tissues (Figure 1C1), and that the percentage of cell expressing RioK1 were 25%, 52.2%, 67.7%, and 87.8% in cancer stage I, II, III, and IV of CRC, respectively (Figure 1C2), revealing that RioK1 expression correlates with CRC malignancy. Importantly, Kaplan–Meier analysis indicated that high levels of RioK1 expression are significantly correlated to overall survival (OS; *P* = 0.003) and disease free survival (DFS; *P* = 0.001) (Figure 1D, Supplementary Table 1). Besides, we also observed an increased expression of RioK1 in gastric cancer (GC) tissues (Figure S1A). Collectively, our data show that the RioK1 expression is frequently upregulated in CRC and GC, and correlated with poor prognosis, suggesting that RioK1 may function as an oncogene in CRC development.

**Figure Legend 1.**
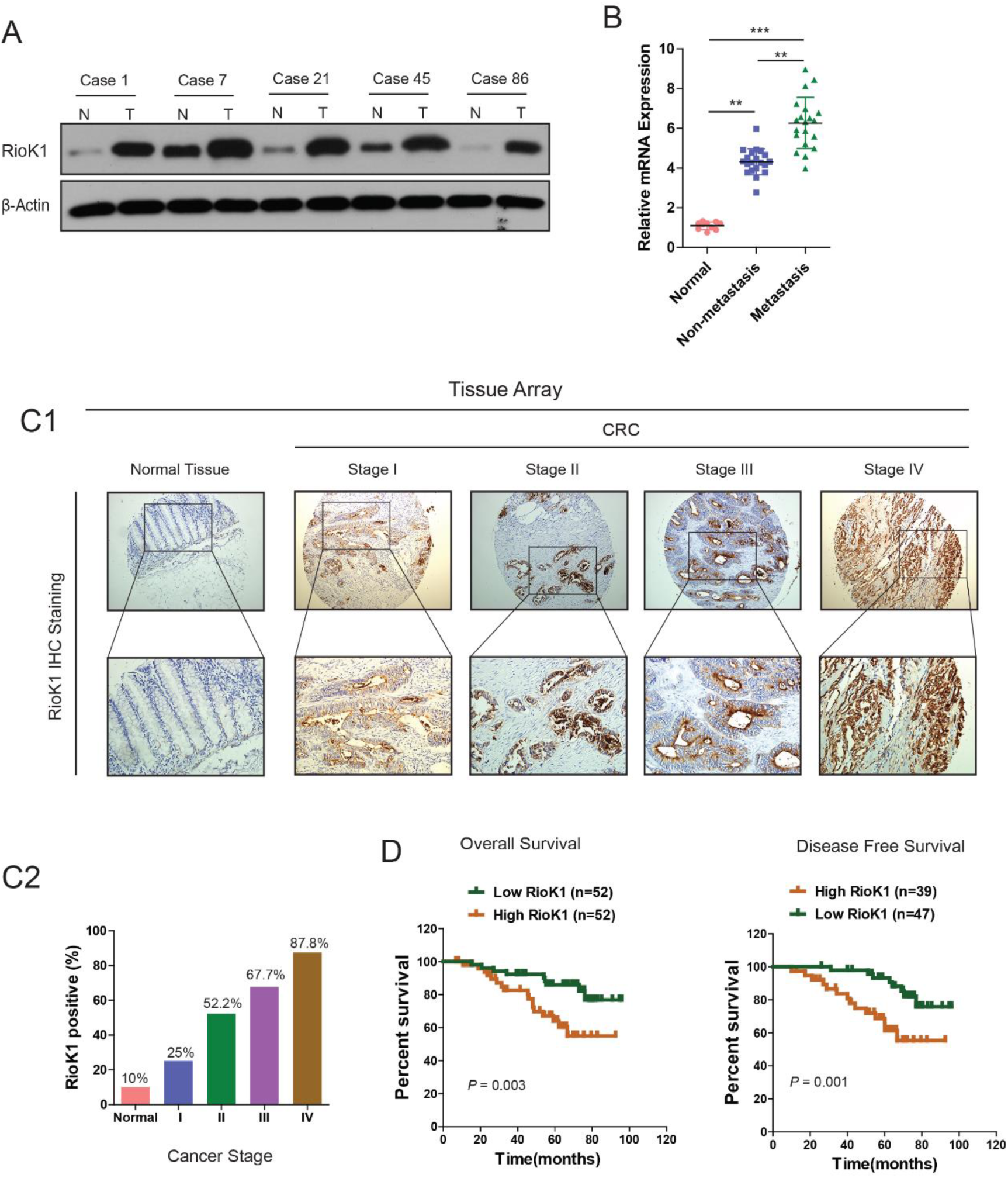
RioK1 is significantly upregulated in CRC and associated with an aggressive and poor survival. (A) RioK1 expression in 5 paired human CRC biopsies and matched normal mucosa analyzed by Western-blot. (B) Comparison of RioK1 mRNA expression level in human CRC tissues (with and without metastasis) and matched normal mucosa. RioK1 mRNA expression was quantified by qPCR and normalized to the matched adjacent normal tissues. (C1) IHC analysis of RioK1 on a tissue micro array of CRC patients (n=110) and healthy adjacent tissue (n=10) using the Allred score.(C2) The IHC signals were scored as 0, 1, 2, and 3; a score ≧ 1+ indicated positive detection. (D) Kaplan meier curves for overall survival and disease free survival of 104 and 86 CRC patients stratified by RioK1 expression respectively.

### RioK1 promotes the proliferation, invasion and metastasis of CRC and GC cells *in vitro* and *in vivo*

Having observed the association of RioK1 expression with poor survival in CRC patients, we set out to functionally characterize the effects of RioK1 on CRC cells. Firstly, we examined the endogenous Riok1 levels of different CRC cell lines and regulated their Riok1 levels using lentivirus-mediated Riok1-specific short hairpin (sh) RNAs or Riok1 plasmid. It was found that the levels of endogenous RioK1 were significantly higher in CRC cell lines than in normal intestinal epithelial cells (IECs) (Figure 2A). Then three Riok1-specific shRNAs (shRiok1) were transfected to silence the endogenous Riok1 expression of CRC cells. According to the western blot analysis, the shRioK1#1 with the highest inhibition rate was selected for further functional experiments (Figure 2B). Both CCK-8 and colony formation assays revealed that down-regulation of RioK1 significantly inhibited the proliferation rate of HCT116 cells compared to negative control (Figure 2C and D). As shown in Figure 2G, both migration and invasion were also obviously inhibited in RioK1 knock-down HCT116 cells. To exclude the possibility of off-target effects, we constructed a shRNA-resistant Riok1 lentiviral vector, RioK1Δ, via site-directed mutagenesis. In HCT116-shRioK1#1 cells, the recovery of RioK1 expression with this shRNA-resistant vector remarkably restored the high proliferative and metastatic ability of the cells (Figure 2C, D, and G). Conversely, overexpression of RioK1 in RKO had the opposite effects on cell viability (Figure 2E and 2F), migration and invasion (Figure 2H). Similar data were observed in SW480 cells transfected with shRioK1#1 and LOVO cells with RioK1 overexpression respectively (data not shown). In summary, the above results illustrated that Riok1 may be critical for the prolieration, invasion and metastasis of CRC cells.

**Figure Legend 2.**
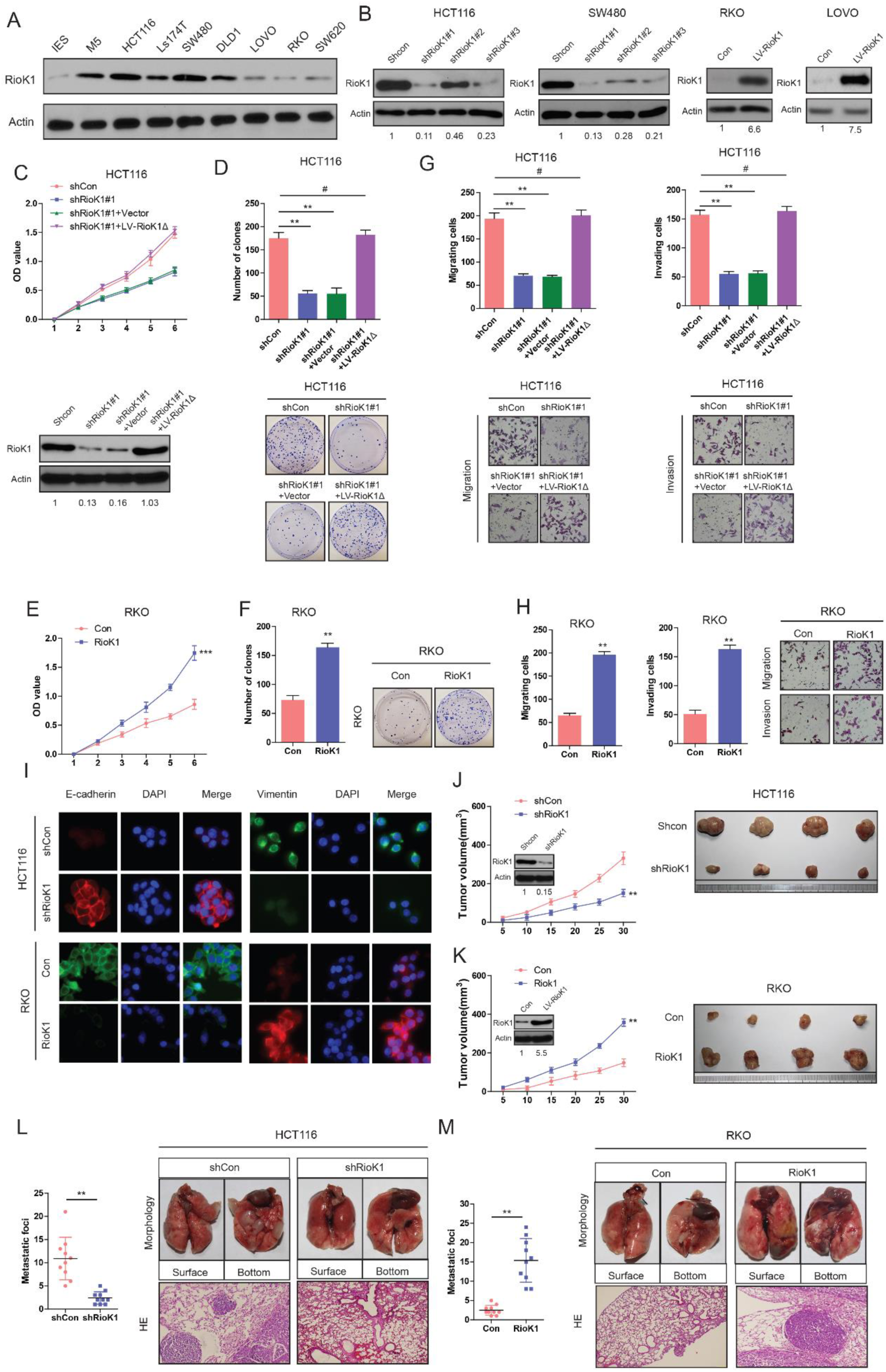
RioK1 promotes growth and metastasis of CRC cells *in vitro* and *in vivo*. (A) The expression levels of RioK1 in 7 CRC cell lines and normal intestinal epithelial cells (IECs) were analyzed by Western blot. (B) Western blot for RioK1 in HCT116 and SW480 cells infected with shRioK1, or control shRNA lentiviral vector, as well as in RKO and LOVO infected with the RioK1-expressing or empty control vector (n = 4). And the intensity of the western blot bands was quantified using NIH ImageJ software. (C) Western blot for RioK1 in the indicated HCT116 cells transfected with shRioK1#1 or the shRNA-resistant expression construct, RioK1Δ. RioK1 expression was recovered in the HCT116-shRioK1 cells transfected with RioK1Δ (lower-panel). And the intensity of the western blot bands was quantified using NIH ImageJ software. RioK1Δ almost restored the proliferative ability of the cells (upper-panel).(D) RioK1 knockdown inhibited proliferation of HCT116 cells determined by CCK-8 and colony formation assays. (E) and (F) Ectopic expression of RioK1 promoted proliferation of RKO cells determined by CCK-8 and colony formation assays. (G) RioK1 knockdown inhibited migration and invasion of HCT116 cells determined by transwell assays. (H) Ectopic expression of RioK1 promoted migration and invasion of RKO cells determined by transwell assays. (I) Representative immunofluorescence images demonstrated that RioK1 level has an effect on the expression of EMT proteins in CRC cells. (J) Growth curve of subcutaneous injection of HCT116/Scramble and HCT116/shRioK1 in NOD/SCID mice (n=6 per group, left panel). NOD/SCID mice were injected subcutaneously into opposite flanks with 1.5 × 10^6^ control cells and RioK1-knocked down HCT116 cells. The mice were sacrificed and the tumors were then removed, weighed and compared (right panel). And knockdown of RioK1 in a xenograft tumor model was analyzed with western blotting, and the intensity of the western blot bands was quantified using NIH ImageJ software. (K) Growth curve of subcutaneous injection of RKO/Vector and RKO/RioK1 in NOD/SCID mice (n=6 per group, left panel). The tumors were then removed, weighed and compared (right panel). Over-expression of RioK1 in a xenograft tumor model was analyzed with western blotting, and the intensity of the western blot bands was quantified using NIH ImageJ software. Effects of RioK1 Knockdown (L) or overexpression (M) on lung metastasis of indicated cells in NOD/SCID mice (*n* = 10 per group): the number of metastatic nodules in the lung (left-panel); representative morphological observation of lung metastases (right-upper panel); and histopathological observation of lung sections (right-lower panel). The results are presented as are means ± SD (*n* = 3 for each panel). Statistical significance was concluded at \**P* < 0.05, \*\**P* < 0.01, \*\*\**P* < 0.001; # represents no statistical significance.

Interestingly, there was a notable change in the expression of epithelial-mesenchymal transition (EMT) proteins in RioK1-overexpressing cells. Cells with relatively high RioK1 expression displayed a mesenchymal-like phenotype, and the expression of vimentin (mesenchymal marker) was enhanced, while RioK1 knockdown strongly upregulated the expression of E-cadherin (epithelial marker) (Figure 2I).

Importantly, an *in vivo* tumor formation assay demonstrated that RioK1 knockdown significantly inhibited the tumorigenesis of CRC cells compared with the control (Figure 2J). Conversely, RioK1 overexpression enhanced the RKO cell growth rate (Figure 2K). Moreover, down-regulation of RioK1 significantly decreased CRC metastatic foci in the lung (Figure 2L) whereas overexpression of RioK1 increased metastatic nodules (Figure 2M). As expected, we observed very similar phenotypes in GC cell lines (Figure S1B, C, D, E, F, G, and H). Taken together, these results indicate that RioK1 promotes growth and metastasis of CRC and GC *in vitro* and *in vivo*.

### RioK1 promotes CRC and GC cell proliferation and migration through PI3K/AKT pathway

Previous studies in Drosophila have indicated that RioK1/2 form a complex with mTOR, which promotes glioblastoma initiation via activating Akt signaling pathway (Read et al., 2013). To identify if the functions of elevated RioK1 expression in CRC and GC cells depend on PI3K/AKT signaling,

CRC and GC cells were incubated with LY294002—an inhibitor of PI3 kinase, or A443654—an inhibitor of Akt kinase. Quantification of the results showed that both proliferation and invasion of RioK1-transfected cells was significantly reduced in LY294002 or A443654 treated cultures (Figure S2A). Meanwhile, treatment of cells with LY294002 or A443654 led to a significant reduction of the levels of Akt phosphorylated at Thr308 and Ser473 (Figure S2B). In parallel, E-cadherin and vimentin, which may account for reversal of epithelial-mesenchymal transition (EMT), were down-regulated and up-regulated respectively in inhibitors-treated cells (Figure S2B), further demonstrating the involvement of PI3K/AKT cascade in the regulation of CRC and GC cells by RioK1.

Conversely, overexpression of constitutively active myristylated Akt (myrAkt), or constitutively high level of active Akt in CRC and GC cells resulting from deficiency in PTEN function (data not shown) abrogated inhibitory effects of RioK1 knockdown on proliferation and invasion when compared with of empty vector, or the domain-negative Akt (DN-Akt) transfected cells (Figure S2C). And the opposite effects on the levels of phosphorylated AKT and EMT-related protein were observed (Figure S2D1 and D2). To confirm the correlation between RioK1 and PI3K/AKT cascade in CRC and GC, we performed immunohistochemistry (IHC) for RioK1 and Akt phosphorylated at Serine-473 on a cohort of CRC and GC specimens. It was found that high RioK1-expressing specimens always showed strong staining for p-Akt-S473 (Figure S2E). These findings further confirmed that PI3K/AKT signaling pathway might contribute to the pro-cancer effects of RioK1 in CRC and GC cells.

### RioK1 interacts with SETD7 *in vitro* and *in vivo*

Given that the activity and biologic role of many kinases are largely dependent on its PTMs (Oliveira and Sauer, 2012), and that previous study implies the involvement of PTMs in regulating the activity and stability of RioK1 (Angermayr et al., 2002), we sought to decide whether RioK1 can be post-translationally modified. To this end, we performed a mass spectrometry analysis of Flag-tagged RioK1 from the cell lysate of HEK293T. Intriguingly, SETD7 was identified as a RioK1-interacting protein from LC-MS/MS analysis (Supplementary Table 2), suggesting the possible involvement of SETD7 in methylation of RioK1.

Firstly, we estimated their physical interaction using an immunoprecipitation (IP) assay in cultured cells. The interaction between GFP-tagged-SETD7 and His-taggged-RioK1 was detected physiologically in RKO cells by using an anti–His antibody followed by Western blot analysis with an anti-GFP antibody (Figure S3A). Again, a reciprocal co-IP assay performed by precipitation with an anti-GFP antibody indicated an interaction between SETD7 and RioK1 (Figure S3A). Co-IP assay also confirmed the binding of SETD7 to RioK1 at endogenous expression levels in RKO cells (Figure S3B). The same co-IP assay was performed in HeLa cells, and a similar result was observed (data not shown), suggesting that the interaction between SETD7 and RioK1 is a universal phenomenon. A GST pull-down assay was performed to further confirm the SETD7 interacted GST-RioK1, but not with GST alone (Figure S3C).

To further map the domains of SETD7 for RioK1 binding, we constructed and purified the Full length (FL) and several fragments of GST-SETD7 and then performed a GST pull-down assay. It was observed that both the N-terminal and middle fragments of SETD7 bound to RioK1 (Figure S3D). Next, the regions of RioK1 for SETD7 binding from the GST pull-down assay were mapped by incubating the FL or fragments of GST-RioK1 with His-SETD7. His-SETD7 mainly interacted with the FL and the 1-120 aa fragment of GST-RioK1, but not with C-terminal fragment of RioK1 and GST alone (Figure S3E). All of the above data indicate that SETD7 directly interacts with RioK1.

### SETD7 monomethylates RioK1 on lysine 411 *in vitro* and *in vivo*

Given that SETD7 has recently been shown to methylate several nonhistone proteins and be preferential for nonhistone proteins, we hypothesized that RioK1 is methylated by SETD7. To test this idea, we firstly examined whether SETD7 interactes with and methylates RioK1 in *Setd7*^-/-^ and *Setd7*^+/+^ MEFs. Consistent with findings described above, co-IP of native RioK1 from *Setd7*^+/+^ MEFs cultured at high density confirmed that SETD7 was in a complex with RioK1 (Figure 3A). When RioK1 was immunoprecipitated and analyzed by immunoblotting with two distinct anti-methyl lysine antibodies (ab23366, recognizing both mono- and dimethylated lysine, and ab7315, primarily recognizing dimethylated lysine), we found that RioK1 is monomethylated in *Setd7*^+/+^ MEFs but not in *Setd7*^−/−^ MEFs (Figure 3A). These results suggest that SETD7 may directly monomethylate RioK1.

**Figure Legend 3.**
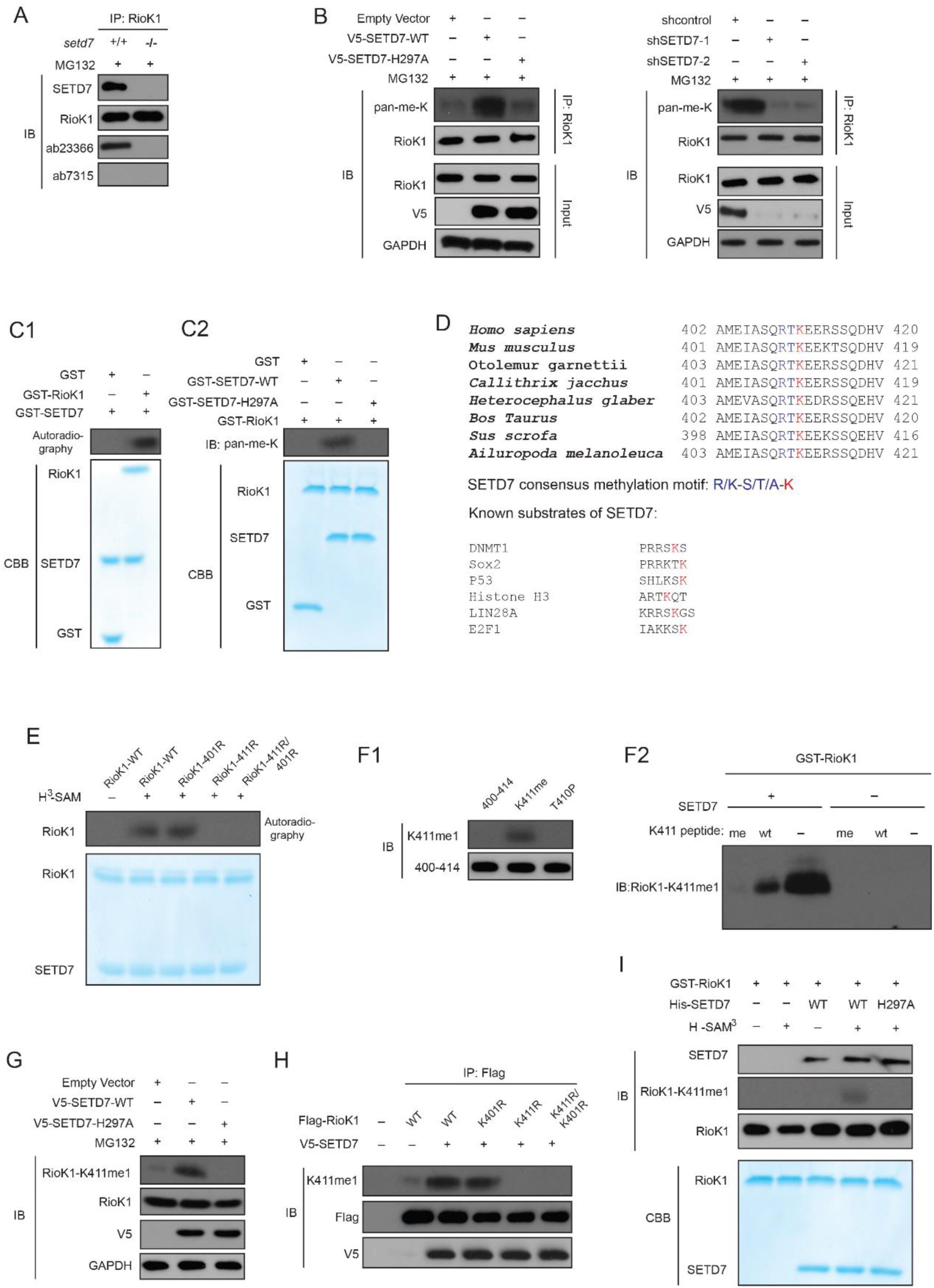
SETD7 monomethylates RioK1 on lysine 411 *in vitro* and *in vivo.* (A) Anti-RioK1 immunoprecipitates from high-density *Setd7*^+/+^ and *Setd7*^−/−^ MEFs were immunoblotted for RioK1, SETD7, methyllysine (mono-, dimethylated) (ab23366), and dimethyl-lysine (ab7315) antibodies, respectively. (B) SETD7 methylates RioK1 *in vivo*. Stably expressed control, SETD7-WT, or SETD7-H297A HEK293T cells were generated. The cell lysates were then precipitated with an anti–RioK1 antibody and probed with an anti–pan-me-K antibody to detect methylation of RioK1 (left panel). Stably expressed shcontrol or sh-SETD7 HEK293T cells were generated. The cell lysates were then precipitated with an anti–RioK1 antibody and probed with an anti–pan-me-K antibody to detect methylation of RioK1 (right panel). (C1) An *in vitro* methylation assay was performed with ^3^H-SAM, recombinant SETD7, and RioK1. Autoradiography and Coomassie brilliant blue R-250 staining (CBB) were used to show methylation and protein levels, respectively. (C2) *In vitro* methylation analysis using indicated proteins. An anti–pan-methyl-lysine antibody (pan-me-K) was used to detect the methylation of RioK1. CBB was used to show the protein levels. (D) Alignment of the consensus amino acid residues adjacent to lysine targeted by SETD7, and identification of a putative SETD7 methylation site in RioK1 (upper panel). The consensus SETD7 recognition sequence (middle panel). The lysines targeted for methylation by SET7 in known substrates are shown (lower panel). In each case, the methylated lysine is shown in red. (E) *In vitro* methylation of various purified RioK1 mutants by SETD7. (F1) An anti-RioK1-K411me1-specific antibody is characterized. RioK1 peptide substrates: 400-414 SKAMEIASQRTKEER, K411me SKAMEIASQRTKmeEER, T410p SKAMEIASQRTpKEER, p: phosphorylation; me: methylation.(F2) GST-RioK1 (1μg) was methylated *in vitro* in the presence or absence of recombinant SETD7 (0.5 μg). The reaction was probed with the anti-RioK1-K411me1 antibody, in the presence of 10 μg/ml competing wild-type (wt) or mono-methylated K411 peptide (me). (G) The effect of ectopically expressed SETD7 and SETD7-H297A on K411me1 of endogenous RioK1 in HEK293T cells. (H) Methylation analysis of Flag-RioK1 and mutants by SETD7 in transfected HEK293T cells. (I) *In vitro* methylation was performed with GST-RioK1, *S*-adenosylmethionine (SAM), recombinant His-SETD7, and His-SET7-H297A. Immunoblotting and CBB were used to show methylation and protein levels, respectively.

To further examine whether the methylation of RioK1 is mediated by SETD7, HEK293T cell lines were transfected with pcDNA (HEK293T-pcDNA), wild-type SETD7 (HEK293T-SETD7-WT), or methylase-deficient mutant SETD7 (HEK293T-SETD7-H297A), and endogenous RioK1 from the cell lines was immunoprecipitated and subjected to Western blot analysis with an anti–pan-methyl-lysine antibody. The protein levels of the methylated RioK1 were significantly increased in the HEK293T-SETD7-WT cell line, but the same phenomenon was not observed in the HEK293T-SETD7-H297A cell line (Figure 3B, left-panel). Conversely, the methylation of RioK1 was almost abolished in the HEK293T-shSETD7 cells compared with the HEK293T-shcontrol cells (Figure 3B, right-panel). Together, these experiments demonstrate that SETD7 methylates RioK1 *in vivo*.

To investigate whether RioK1 is methylated directly by SETD7, an *in vitro* methylation assay was performed by incubating SETD7 with ^3^H-SAM. As shown in Figure 3C1, GST-RioK1 was methylated by SETD7, but not by a catalytic mutant SETD7-H297A (Figure 3C2), which confirms that SETD7 methylates RioK1 *in vitro*.

Next, we sought to identify the SETD7-methylated residue(s) in RioK1. The amino acid sequence of RioK1 was compared with the sequences surrounding the SETD7 methylation sites of several known substrates (Figure 3D). It was found that the conserved lysine residue at position 411 and its neighboring residues of RioK1 strongly conform to the consensus SETD7 recognition sequence designated by K/R-S/T/A-K motif (in which the methylation lysine site is underlined). When we aligned the full amino acid sequences of RioK1 proteins of various species, it was observed that the identified RioK1 methylation sequence was evolutionarily conserved (Figure 3D). Moreover, K411R mutation almost completely abolished RioK1 methylation by SETD7 *in vitro*, whereas other mutation such as K401R had no effect (Figure 3E). Together, our data suggest that K411 is most likely the major, if not the sole, site within RioK1 that is methylated by SETD7.

To easily find whether methylation of RioK1 occurred in cells, we prepared a modification-specific antibody, anti-RioK1-K411me1, which specifically recognized monomethylated RioK1-K411 but not the unmethylated peptide (Figure 3F1 and Figure S4). This antibody detected methylated GST-RioK1 after treatment with SETD7, with minimal reactivity against untreated GST-RioK1 (Figure 3F2). Further, the unmodified RioK1 peptide poorly prevented the anti-RioK1-K411me1 antibody binding to methylated GST-RioK1, whereas the methylated peptide prevented binding of the anti-RioK1-K411me1 antibody (Figure 3F2).

Using this RioK1-K411me1-specific antibody, it was observed that ectopic expression of V5-SETD7 in the HEK293T cells led to substantially increased endogenous RioK1-K411me1, while the enzymatic deficient SETD7-H297A mutant reduced the level of RioK1-K411me1 (Figure 3G). Furthermore, V5-SETD7 catalyzed K411me1 for Flag-RioK1-WT and Flag-RioK1-K401R mutant, but not K411R mutant (Figure 3H). The specificity of anti-RioK1-K411me1 toward RioK1 was further confirmed using an *in vitro* cell-free methylation reaction with purified recombinant GST-RioK1 and His-SETD7 in the presence of *S*-adenosylmethionine (Figure 3I). Collectively, these results demonstrate that SETD7 methylates RioK1 at K411 both *in vitro* and *in vivo*.

### Methylation of RioK1 by SETD7 significantly decreases its stability

Lysine methylation is usually linked to the stability of nonhistone, including RelA, DNMT1, Sox2, p53, ERα, RelA, and FoxO3 (Couture et al., 2006; Wang et al., 2013; Estève et al., 2011; Li et al., 2008). Next, we tested whether the methylation of RioK1 by SETD7 affects its stability in cells. It was found that SETD7 overexpression decreased the expression of RIOK-WT but not of RioK1-K411R in HEK293T cells (Figure 4A), and mRNA levels of RioK1 was not changed (Figure 4B), suggesting that the methylation of RioK1 may disrupt its stability. In HEK293T-shcontrol cells, the half-life of endogenous RioK1 was 4–8 h, while in HEK293T-shSETD7-1 cells it was dramatically prolonged (Figure 4C and D). To evaluate whether this change in the stability of RioK1 depends on its methylation at K411, the effect of SETD7 on the half-life of RioK1-WT and –K411R was investigated. Consistently, the half-life of RioK1-WT was decreased from 4–8 h to < 4 h when RioK1-WT was coexpressed with SETD7 (Figure 4E and F). Conversely, RioK1-K411R was more resistant to SETD7-induced degradation, with a much longer half-life (> 8 h) (Figure 4E and F). Collectively, our data demonstrate that the methylation of RioK1 at K411 by SETD7 reduces its stability.

**Figure Legend 4.**
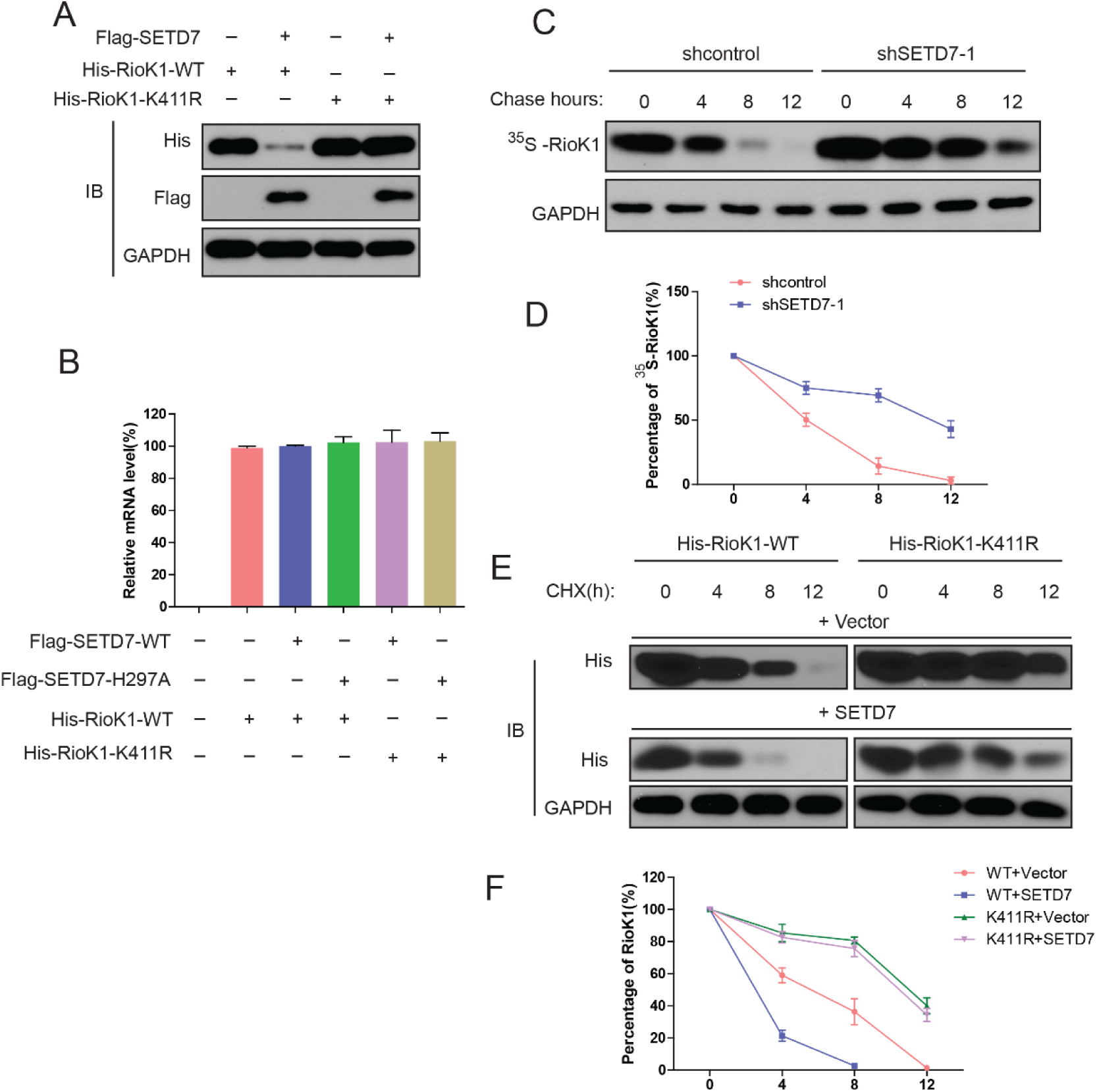
SET7/9 significantly reduces the stability of RioK1. (A) His-RioK1-WT or His-RioK1-K411R was cotransfected into HEK293T cells for 48 h with or without Flag-SET7/9. Cell lysates were examined by western blot with indicated antibodies. (B) His-RioK1-WT was cotransfected with or without Flag-SET7/9 or mutant Flag-SET7/9-H297A into HEK293T cells for 48 h. Real-time PCR was used to detect the mRNA levels of RioK1. (C) HEK293T-shcontrol and HEK293T-sh-SET7/9-1 cells were metabolically labeled with [^35^S]-methionine, as indicated, and followed by chasing with standard medium. Total lysates were examined with anti-RioK1 antibody. (D) Quantification analysis of endogenous RioK1 levels in (C). (E) HEK293T cells were transfected with indicated plasmids, and 24 h later, treated with cycloheximide (CHX), and probed with anti-His antibody. (F) Quantification analysis of the results in (E).

### RioK1 demethylation by LSD1 increases RioK1 stability

Our mass spectrometric analysis of the RioK1 immunocomplex from HEK293T cells uncovered that lysine-specific demethylase 1 (LSD1), also known as KDM1A, one component of a BRAF–HDAC complex (BHC) histone deacetylase complex (Iwase et al., 2004), was present in the immunocomplex (Supplementary Table 1). Given that LSD1 can demethylate non-histone proteins p53 and Dnmt1 (Jin et al., 2013; Nicholson et al., 2009), we hypothesized that LSD1 may reverse SETD7-induced RioK1 methylation. Indeed, endogenous RioK1 was observed to be present in LSD1 immunocomplex, and endogenous LSD1 was also present in the RioK1 immunocomplex in 293T cells (Figure 5A). To exclude the possibility of a false-positive interaction resulting from cross-reactivity of antibodies, we demonstrated that Flag-RioK1 physically associated with HA-LSD1 in 293T cells in the reciprocal coimmunoprecipitation assays (Figure 5B). When GST-RioK1 and His-LSD1 were mixed together, it was observed that GST-RioK1 specially pulled down His-LSD1 (Figure 5C), indicating that LSD1 directly binds RioK1.

**Figure Legend 5.**
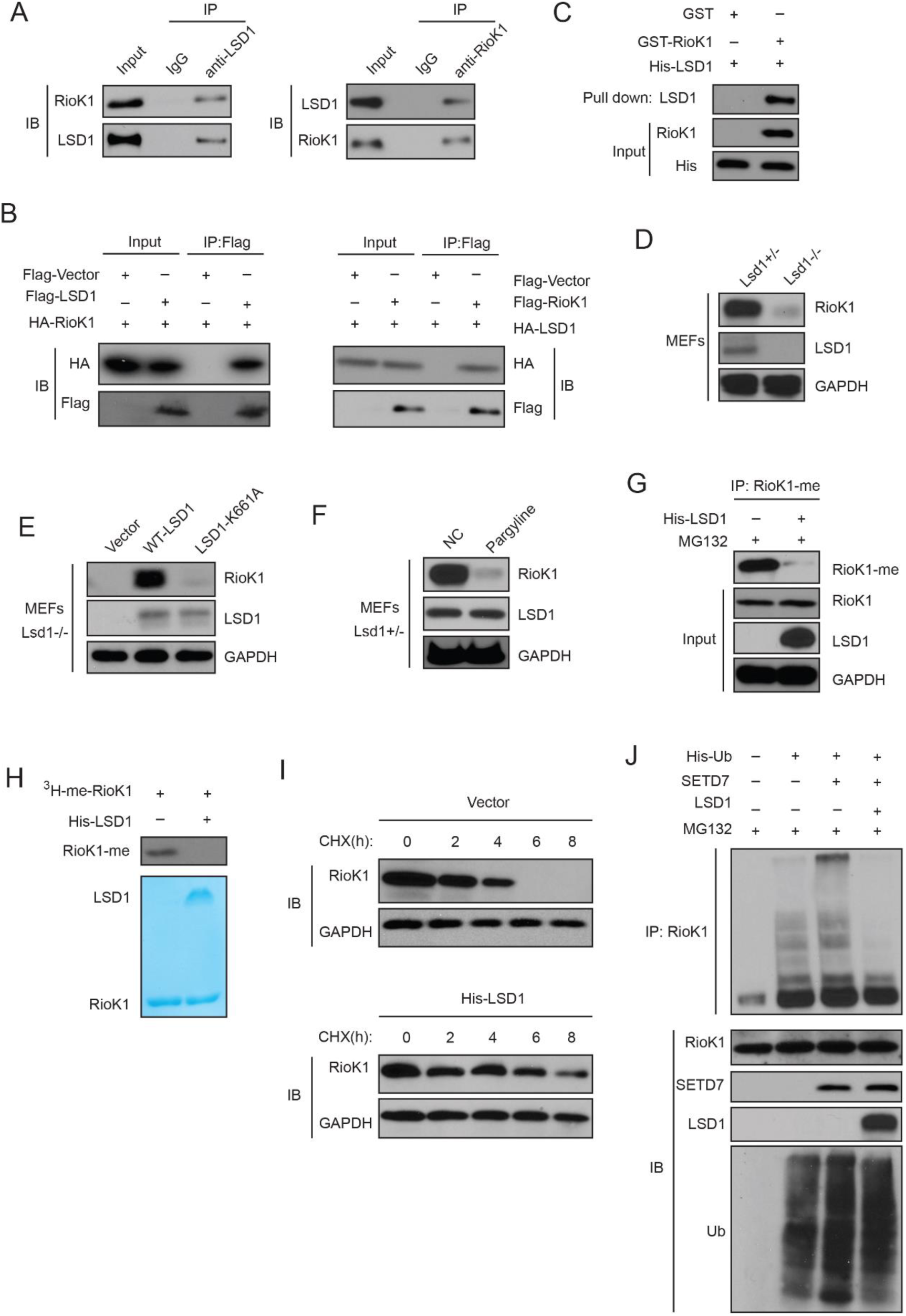
SET7/9-mediated methylation of RioK1 is reversed by LSD1, which increases RioK1 stability. (A) Endogenous RioK1 co-immunoprecipitated reciprocally with endogenous LSD1 from HEK293T cells and subjected to immunoprecipitation followed by immunoblotting with indicated antibodies. (B) HA-RioK1 co-immunoprecipitated reciprocally with Flag-LSD1. Total cell lysates were extracted from HEK 293T cells transiently cotransfected with expression constructs as indicated and probed with antibodies as indicated. (C) RioK1 directly interacted with LSD1. Bacterially produced GST-RioK1 was used to pull down bacterially produced His-LSD1 *in vitro*. (D) RioK1 protein levels in *Lsd1^+/−^* and *Lsd1^−/−^* MEFs were compared. (E) *Lsd1^−/−^* MEFs were reconstituted with either WT or a catalytically inactive Mutant (K661A) of LSD1. RioK1 protein levels were monitored. (F) *Lsd1^+/−^* MEFs were pretreated with pargyline for 12 h, RioK1 protein levels were evaluated. (G) Overexpressing LSD1 with MG132 treatment significantly decreases RioK1 methylation in HEK293T cells by. (H) LSD1 reverses SET7/9-dependent RioK1 methylation. *In vitro* demethylation assays using purified His-LSD1 were performed and RioK1 methylation was detected by autoradiography.(I) HEK293T cells transfected with indicated plasmids were treated with CHX (20 μg /ml), collected at the indicated times, and analysed by western blot. (J) overexpressing LSD1 with MG132 treatment from HEK293T cells co-transfected with the indicated plasmids. And cell lysates were subjected to pull-down with Ni^2+^-NTA beads. RioK1 ubiquitination was assessed by anti-RioK1 antibody in the presence of MG132.

As expected, compared with the protein levels of RioK1 in *Lsd1*^+/−^ MEFs, those in *Lsd1*^−/−^ MEFs were found to be greatly decreased (Figure 5D). Reconstitution experiments with either LSD1 WT or an enzymatically inactive LSD1-K661A mutant in *Lsd1*^−/−^ MEFs further indicated that RioK1 protein levels were restored in only LSD1 WT-reconstituted cells but not in LSD1-K661A mutant-reconstituted cells (Figure 5E). Moreover, treatment of pargyline, a LSD1 inhibitor which blocks LSD1 enzymatic activity, promoted RioK1 destabilization (Figure 5F).

Then we tested whether LSD1 is responsible for RioK1 demethylation. Indeed, LSD1 led to RioK1 demethylation both *in vivo* and *in vitro* (Figure 5G and H). Moreover, treatment of cycloheximide (CHX) showed that SETD7 overexpression significantly decreased the half-life of endogenous RioK1, while ectopic LSD1 expression increased it (Figure 5I). To confirm the involvement of 26S proteasome-dependent degradation pathway in the stabilty of RioK1 protein, we performed RioK1 ubiquitination assay with SETD7 or LSD1 in the presence of MG132. Unsurprisingly, SETD7 significantly increased RioK1 ubiquitination whereas the overexpression of LSD1 almost completely abolished the increase in RioK1 ubiquitination (Figure 5J). Collectively, our results suggest that LSD1-dependent demethylation of RioK1 significantly stabilizes RioK1 proteins by reversing SETD7-mediated RioK1 methylation-dependent degradation by 26S proteasomes.

### SETD7 promotes RioK1 ubiquitination and degradation through the F Box Protein FBXO6

Next, we set out to identify the E3 ubiquitin ligase that mediates K411me-dependent RioK1 ubiquitination and degradation. Since both FBXW7 and FBXO6 were present in the RioK1 immunocomplex (supplementary Table 1), we focused our attention on the Cul1-containing (Skp1-Cul1-F box) SCF E3 ligase complex. We found that knockdown of FBXO6, but not FBXW7, by two different shRNAs increased not only levels of RioK1 proteins (Figure 6A), but also the stability of RioK1 protein in cells treated with CHX (Figure 6B).

**Figure Legend 6.**
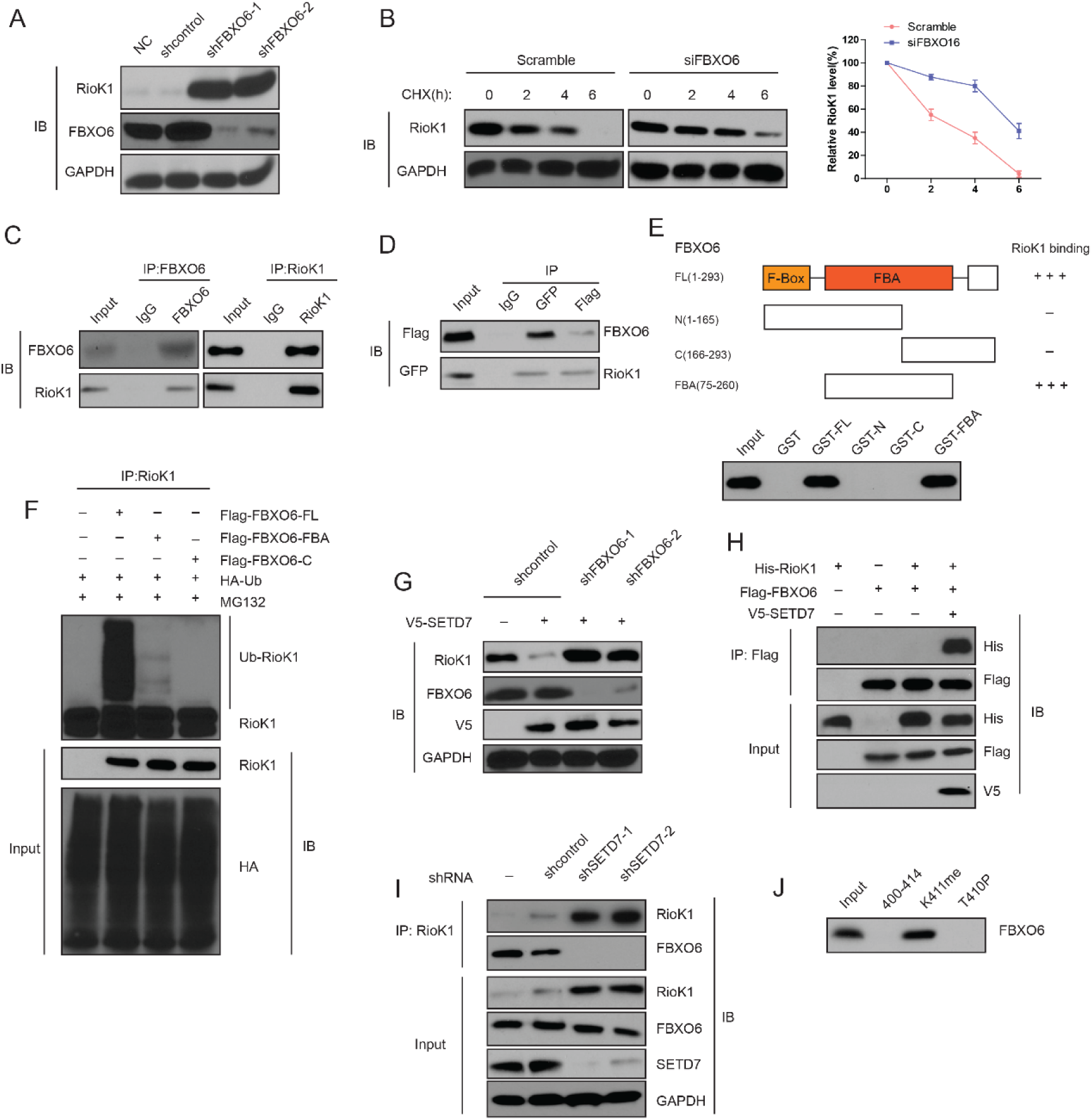
K411 methylation of RioK1 by SETD7 promotes the interaction between FBXO6 and RioK1. (A) The effect of FBXO6 knockdown on the levels of RioK1 proteins in HEK293T cells. (B) HEK293T cells were transfected with the indicated siRNAs, and after 48 hr, treated with 160 μM CHX, and representative RioK1 expression result is shown in the left panel. Right panel indicates quantitation of the RioK1 blots. (C) HEK293T cells were extracted and immunoprecipitated with an anti–FBXO6 (left) or anti-RioK1 (right) antibody. Western blot analysis was performed with indicated antibodies. (D) Whole-cell lysates of HEK293T cells transfected with GFP-FBXO6 and Flag-RioK1 were precipitated with an anti-GFP or anti-Flag antibody, and the interactive components were analyzed by Western blot. (E) Binding of several different domains of human FBXO6 to RioK1. Numbers represent the amino acid residues in human FBXO6. FL, N, C, and FBA represent the full-length, amino terminus, carboxyl terminus, and the FBA domain only of FBXO6, respectively. The extent of the interaction between RioK1 and FBXO6 domains is indicated by the number of plus signs (+). (F) HEK293T cells were transfected with HA-RIOK, Myc-tagged FBXO6 FL or mutants with His-ubiquitin for 48 hr. Cells were lysed and blotted with indicatd antibodies. (G) Knockdown of FBXO6 blocked V5-SETD7-induced RioK1 degradation. (H) The interaction between Flag-FBXO6 and HA-RioK1 was dependent on SETD7. (I) SETD7 knockdown disrupted the interaction between endogenous FBXO6 and RioK1 in HEK293T cells. (J) The binding of FBXO6 to unmodified, T410p, or K411me RioK1 aa 400–414 peptides was analyzed using *in vitro* peptide pull-down assay.

Next, we estimated their physical interaction using an immunoprecipitation assay in cultured cells. Endogenous RioK1 was immunoprecipitated from 293T cells, and subsequent immunoblotting revealed that FBXO6 was immunoprecipitated with RioK1 (Figure 6C). Reverse immunoprecipitation of FBXO6 confirmed the identified interaction with RioK1 (Figure 6C). The interaction between RioK1 and FBXO6 was also obvious when a co-IP assay was performed in 293T cells with the overexpression of GFP-SETD7 and Flag-RioK1 (Figure 6D). To further define the RioK1-FBXO6 interaction, we generated several polypeptides corresponding to different domains of human FBXO6. It was observed that the F box associated (FBA) domain mediates the binding of FBXO6 to RioK1 (Figure 6E).

To examine the role of FBXO6 in RioK1 ubiquitination, we transfected 293T cells with HA-RioK1, His-ubiquitin, and different domains of human FBXO6. Interestingly, poly-ubiquitination of HA-RioK1 was readily observed in the cells with FBXO6 full length (FL), but was significantly less noticeable in cells cotransfected with the FBXO6-derived FBA (Figure 6F), demonstrating that only the FBXO6 FL protein supported ubiquitination of RioK1 *in vitro*. Together, these data suggest that FBXO6 is a ubiquitination E3 ligase of RioK1.

We next investigated whether FBXO6 mediates SETD7-induced RioK1 degradation. Western blot analysis confirmed that knockdown of FBXO6 by two different shRNAs blocked RioK1 degradation induced by SETD7 (Figure 6G), suggest that FBXO6 is required for SETD7-induced RioK1 degradation.

Next question was how SETD7-induced RioK1-K411me is functionally linked to FBXO6. We did not observe the binding of FBXO6 to SETD7 (data not shown). When V5-SETD7 was co-expressed, Flag-FBXO6 was found to associate with HA-RioK1 (Figure 6H). However, Flag-FBXO6 did not bind to HA-RioK1 when V5-SETD7-H297A was co-expressed, and Flag-FBXO6 failed to coimmunoprecipitate with RioK1-K411R mutant with or without co-expressed V5-SETD7 (data not shown). Furthermore, Flag-FBXO6 was found to interact with RioK1 in the HEK293T cellular extracts, however, upon knockdown of SETD7, this interaction was abolished (Figure 6I). Finally, *in vitro* peptide pull-down assay indicated that the full-length FBXO6 specifically bound to the K411me peptide, not the peptide phosphorylated at T410 (T410p) (Figure 6J). Collectively, our data suggest that SETD7 promotes the interaction between FBXO6 and RioK1 through its ability to methylate RioK1 at K411.

### Casein kinase 2 (CK2) phosphorylates RioK1 both *in vitro* and *in vivo*

Given that the full-length FBXO6 only bound specifically to the K411me peptide, not the peptide phosphorylated at T410 (T410p), and especially threonine residue at 410 is exactly located within the consensus sequences of SETD7, we reasoned that T410 phosphorylation of RioK1 might influence the biological functions of SETD7 –mediated K411 methylation of RioK1. To test this idea, we firstly aimed to identify the kinase responsible for the phosphorylation of threonine residue at 410 (T410) on RioK1. The residues surrounding T410 of RioK11 (RTKEER), with acidic residues at +1 and +4, conform to a putative protein kinase CK2 phospho-motif (S/TXXE/D), which makes T410 an optimal site for phosphorylation by CK2 (Figure 7A). Thus, we investigated whether CK2 mediated the phosphorylation of RioK1 at T410 and what functional relevance this modification had in various cell types (including CRC cells).

**Figure Legend 7.**
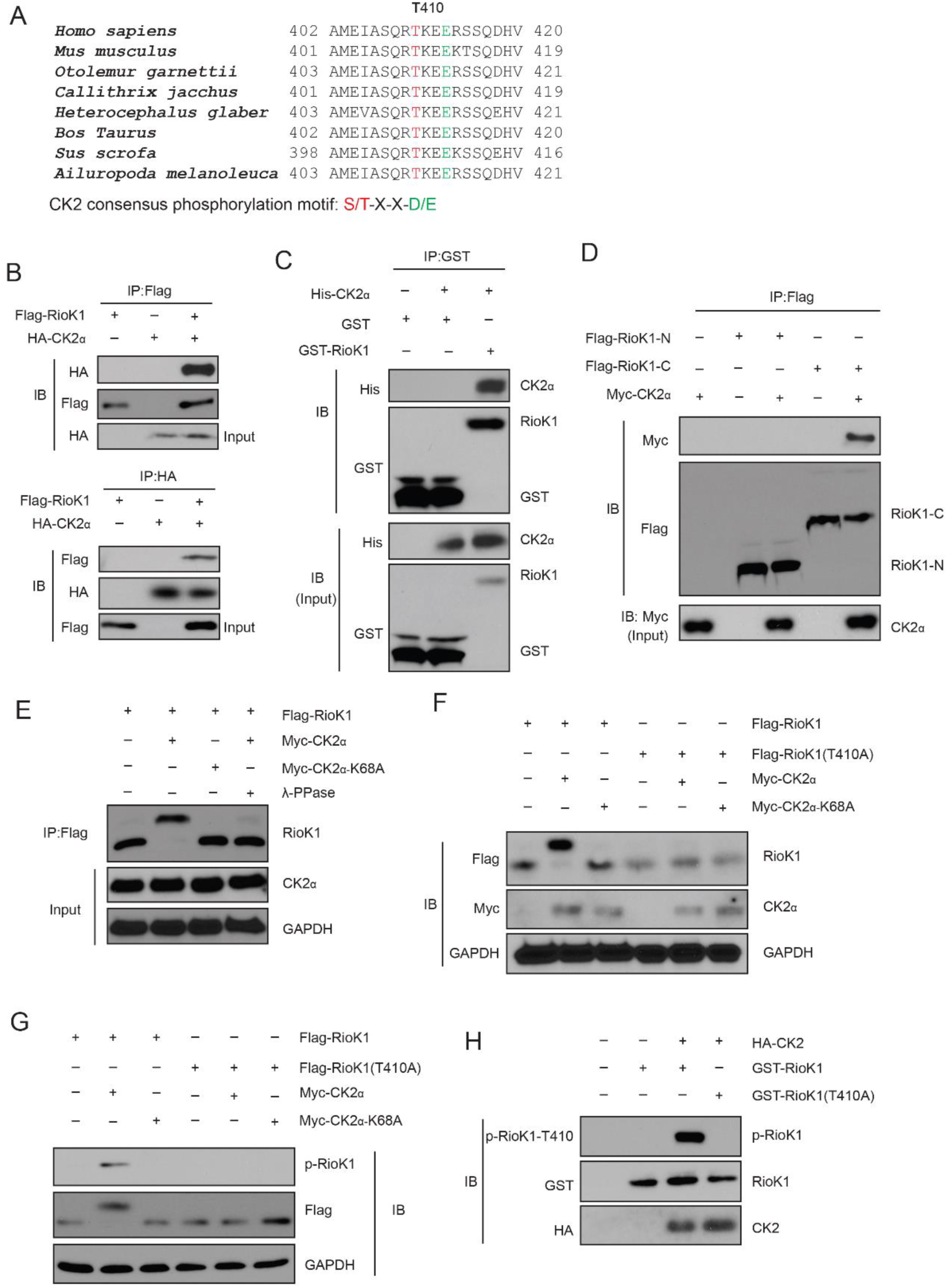
CK2 phosphorylates RioK1 at T410 both in vitro and in vivo. (A) RioK1 consensus phosphorylation sites (shown in red) corresponding to the CK2 consensus motifs S/TXXE/D are presented. These consensus phosphorylation sites is exactly located within the consensus sequences of SETD7. (B) FLAG-RioK1 was present in the HA-CK2 immunocomplex. Total cell lysates were extracted from 293T cells transiently co-transfected with FLAG-RioK1 and HA-Vec and HA-CK2α subjected to immunoprecipitation with an anti-HA antibody followed by immunoblotting with indicated antibodies. (C) *In vitro* glutathione *S*-transferase (GST)-precipitation assay of no CK2α, or purified His-tagged CK2α combined with GST alone or GST-RioK1. (D) Immunoprecipitation and immunoblot analysis of lysates of HEK293T cells expressing no plasmid (−) or plasmid encoding the Flag-tagged amino terminus (amino acids 1–242) of RioK1 (Flag–RioK1-N) or carboxyl terminus (amino acids 243–568) of RioK1 (Flag–RioK1-C), plus Myc-tagged CK2α, probed with anti-Myc and/or anti-Flag. (E) Lysates of HEK293T cells expressing Flag-tagged wild-type RioK1 and Myc-tagged wild-type CK2α or CK2α-K68A; far right (λ-PPase), was analyzed using Western blot. GAPDH serves as a loading control throughout. (F) Immunoblot analysis of RioK1 and CK2α in lysates of HEK293T cells expressing Flag-tagged wild-type RioK1 or RioK1 (T410A) and Myc-tagged wild-type CK2α or CK2α-K68A. (G) Immunoblot analysis of phosphorylated p-RioK1-T410 and total RioK1 and CK2 in lysates of HEK293T cells. (H) *In vitro* kinase assay of purified recombinant GST-tagged RioK1 or RioK1-T410A with HA-tagged CK2.

CK2 is a constitutively active serine/threonine kinase that is ubiquitously expressed (Bollacke et al., 2016). Its tetrameric holoenzyme is typically composed of two catalytic subunits (CK2α and CK2α’) and two regulatory β subunits (Bollacke et al., 2016). As expected, we confirmed that both HA-tagged CK2α and CK2α’ (data not shown) physically associated with FLAG-tagged RioK1 (Figure 7B). We further found that endogenous RioK1 in 293T cells coimmunoprecipitated with endogenous CK2α, and with CK2α’ to a lesser extent, and that endogenous CK2α bind to endogenous RioK1 (data not shown). Purified GST-CK2α was able to pull down bacterially produced His-RioK1 (Figure 7C), indicating that the interaction between RioK1 and CK2 is direct. We further found that the carboxy-terminal domain of RioK1 is required for its binding to CK2 (Figure 7D).

We next investigated the effect of CK2 on RioK1. When co-expressed with CK2α, but not CK2α-K68A, a kinase-inactive mutant of CK2α (Lebrin et al., 2016), RioK1 was found to migrate more slowly only in the presence of CK2α than did RioK1 on its own by SDS-PAGE (Figure 7E), suggesting that this differential migration was the result of phosphorylation. Threonine 410 residue is conserved in RioK1 from various species, and is homologous to the CK2-phosphorylated substrate motif (Figure 7A). Substitution of the threonine at position 410 with alanine (T410A) abolished the CK2-induced slower migration of RioK1 by SDS-PAGE (Figure 8F). To further study the phosphorylation of RioK1 by CK2, we generated a rabbit polyclonal antibody to RioK1 phosphorylated at T410 (p-T410) by immunizing rabbits with the RioK1 peptide (data not shown). Co-expression of wild-type CK2α, not CK2α-K68A, with RioK1 induced phosphorylation of RioK1 at T410, and the T410A substitution in RioK1 eliminated the phosphorylation of RioK1 by CK2 (Figure 7G). In an *in vitro* kinase assay, purified RioK1 was phosphorylated by CK2, but mutant RioK1 (T410A) did not (Figure 7H). Collectively, our results suggest that CK2 directly phosphorylated RioK1 at T410.

**Figure Legend 8.**
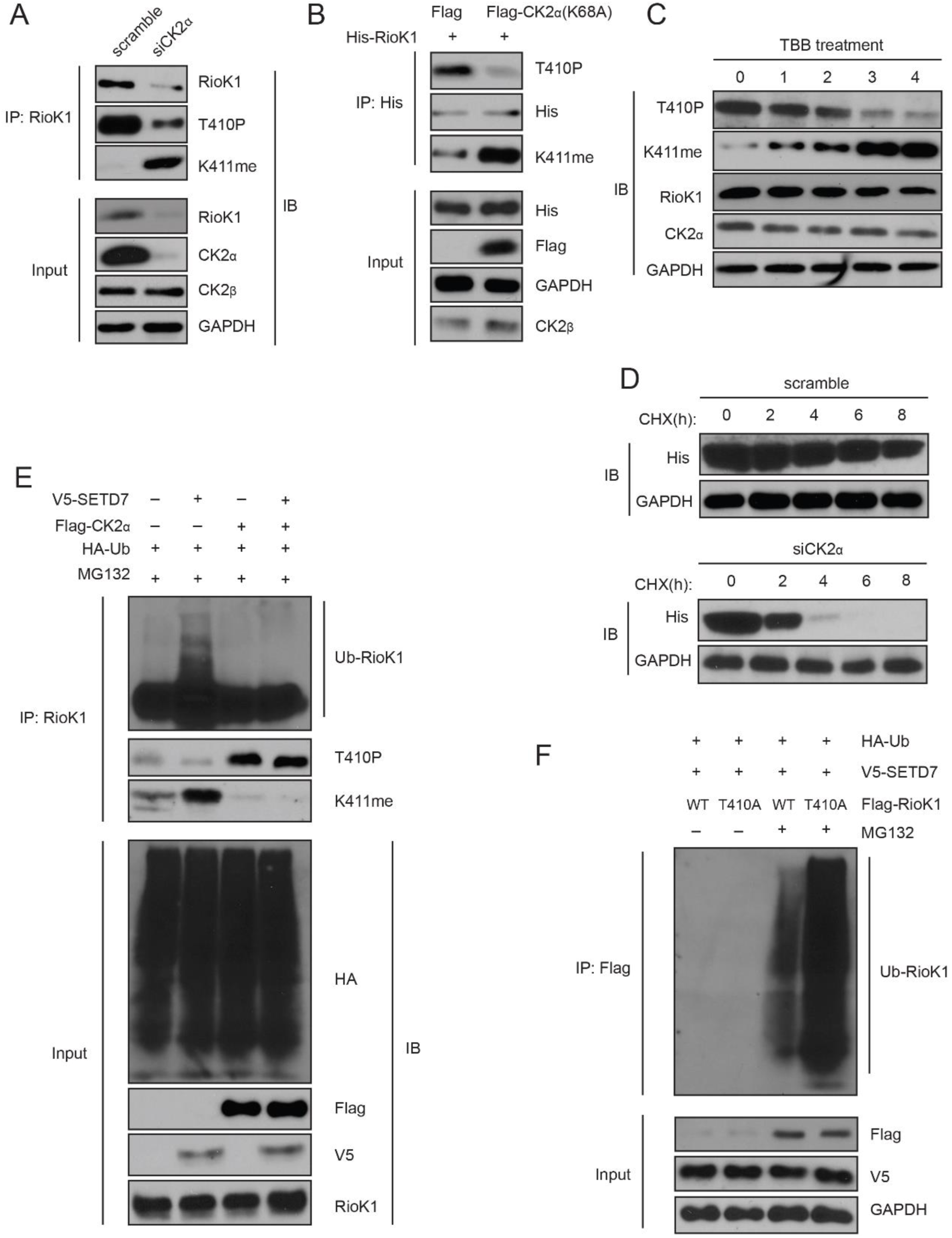
T410 phosphorylation of RioK1 antagonists SETD7-mediated K411 methylation, which stabilizes RioK1 in CRC and GC cells. (A) The effect of knockdown of CK2α in HCT116 cells on the levels of RioK1 proteins, T410p, and K411me. (B) The effect of the CK2α-K68A mutant on the levels of RioK1 proteins, T410p, and K411me. (C) The effect of the CK2 kinase inhibitor TBB on the levels of RioK1 proteins, T410p, and K411me. (D) The effect of CK2 knockdown in HCT116 cells on the RioK1 protein stability. (E) CK2 inhibits K411me by SETD7 and its consequent RioK1 ubiquitination in RKO cells. Indicated cells were cotransfected with HA-Ub and treated with MG132 for 12 hr. (F) Wild-type RioK1 or the RioK1-K411me1 mutant was overexpressed in cells, along with HA-ubiquitin and V5-SET7, in the presence or absence of the proteasome inhibitor MG132. Cell lysates were analyzed with indicated antibodies.

### RioK1 phosphorylation at T410 by CK2 blocks methylation at K411 by SETD7 and stabilizes RioK1

Given the close proximity of T410 and K411 residues, we analyzed whether T410p by CK2 could affect K411me by SETD7. Indeed, knockdown of CK2α in HCT116 cells led to a substantial reduction of RioK1 protein and p-T410, and a significant increase of RioK1 K411me (Figure 8A). Consistently, in the presence of CK2α-K68A, both RioK1-p-T410 and total RioK1 level were less abundant, and RioK1-K411me1 levels increased (Figure 8B).

To confirm that RioK1 stability is CK2 dependent, HCT116 cells were treated with the CK2 inhibitor TBB (Leung et al., 2015). RioK1-p-T410 levels substantially decreased within 3 h of the treatment, while RioK1-K411me proportionally increased (Figure 8C). Along with the gradual increase of RioK1-K411me, more degradation of RioK1 was observed, causing a decrease in total RioK1 levels, however, the total level of CK2α remained similar (Figure 8C). Additionally, knockdown of CK2α significantly reduced half-life of RioK1 (6–8hr in control versus less than 4hr) in HCT116 cells (Figure 8D). These data suggest that CK2-induced phosphorylation of RioK1 at T410 might stabilize RioK1 by inhibiting K411me by SETD7.

Next, we investigated how CK2 influences RioK1 ubiquitination. While overexpression of SETD7 in RKO cells led to increased K411me and RioK1 ubiquitination, overexpression of CK2α significantly increased p-T410 and inhibited RioK1 ubiquitination (Figure 8E). Importantly, co-expression of CK2α with SETD7 only significantly led to an increase in p-T410, but not K411me, suggesting that p-T410 by CK2 predominates over and inhibits K411me by SETD7 (Figure 8E). Consistent with this idea, CK2α overexpression blocked SETD7-induced RioK1 ubiquitination (Figure 8E). We observed similar results when the experiments were performed in MKN45 cells (data not shown). Therefore, compared to the methylated RioK1, the phosphorylated species is less likely to be degraded.

Next, we tested whether the T410A mutation promoted ubiquitin-induced degradation of RioK1. Unsurprisingly, in the presence of MG132, transfection with V5-SETD7 and Flag-RioK1-WT led to less ubiquitination than transfection with V5-SET7 and Flag-RioK1-T410A (Figure 8F), suggesting that the T410A mutation promotes RioK1 methylation which is a substrate for ubiquitinylation.

Collectively, our data demonstrate that p-T410 by CK2 blocks K411me by SETD7, which protects RioK1 from ubiquitination and degradation.

### SETD7 –mediated methylation of RioK1 negatively regulates CRC growth and metastasis, which is reversed by the CK2-induced phosphorylation of RioK1

To explore the functional significance of both methylation and phosphorylation of RioK1, we evaluated the impact of an unmethylatable mutation K411R, a non-phosphorylatable mutation T410A, and an unmethylatable plus phosphomimetic mutation (K411R/T410E) of RioK1 on CRC cell proliferation, migration, invasion, tumor growth and metastasis. MTT assays showed increased growth in cells stably expressing RioK1-K411R and RioK1-K411R/T410E, but attenuated growth in cells with RioK1-T410A compared to cells with RioK1-WT expression (Figure 9A). The ability to form colonies was dramatically increased in stable transfectants with the expression of RioK1-K411R and RioK1-K411R/T410E, compared to RioK1-WT (Figure 9B). In agreement with these *in vitro* growth assays, RKO cells with stable expression of RioK1-K411R and RioK1-K411R/T410E showed greater tumor growth than cells with RioK1-WT in a xenograft tumor model, while the stable expression of RioK1-T410A stable obviously retarded CRC growth (Figure 10C and D).

**Figure Legend 9.**
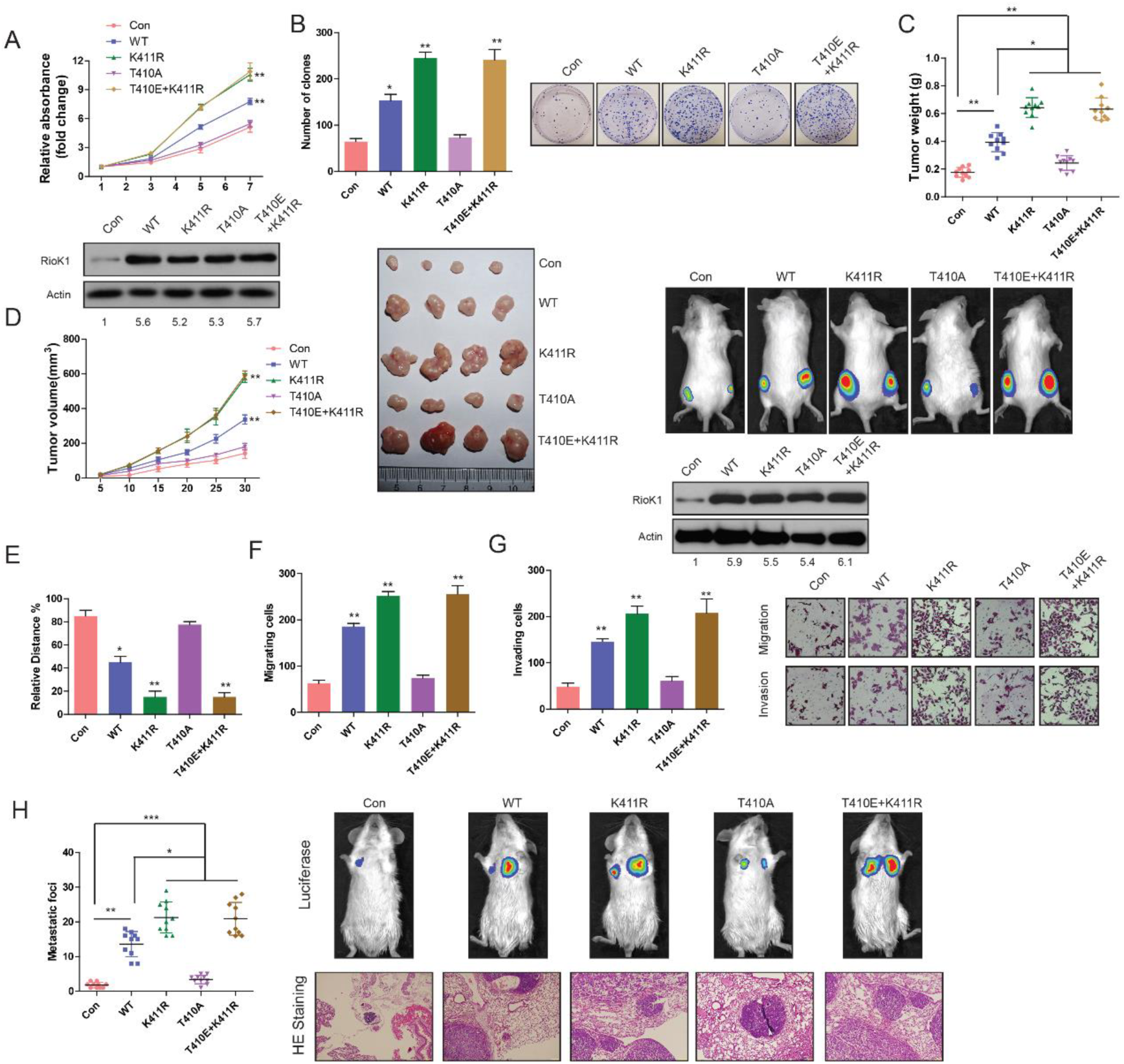
Methylation of RioK1 at K411 and phosphorylation of RioK1 at T410 oppositely regulated the CRC growth and metastasis. (A) MTT assays in RKO cells expressing indicated proteins (upper panel). Western blotting was used to detect expression of RioK1 in RKO cells using anti-RioK1 antibody (lower panel), and the intensity of the western blot bands was quantified using NIH ImageJ software. (B) Photography and statistic results of colony formation assays in RKO cells expressing indicated proteins. (C and D) Statistic, tumor growth curves, photography results and representative bioluminescence images of tumor size in RKO cells expressing indicated proteins. Over-expression of RioK1 in a xenograft tumor model was analyzed with western blotting, and the intensity of the western blot bands was quantified using NIH ImageJ software. (EG) Migration ability evaluated by wound healing assay (E) and transwell migration assay (F). Invasiveness evaluated by matrigel invasion assay (G) in RKO cells expressing indicated proteins. (H) Statistic results, HE staining and representative bioluminescence images of lung metastasis in RKO cells expressing indicated proteins.

To assess cell migration changes, wound healing and transwell migration assays were performed in CRC cells. As shown in Figure 10E and F, CRC cells with the expression of RioK1-K411R and RioK1-K411R/T410E had enhanced migration ability compared to RioK1-WT. Similarly, the invasion ability of CRC cells stably expressing RioK1K411R and RioK1-K411R/T410E was also substantially increased (Figure 9G). In agreement with these *in vitro* migration assays, stable expression of RioK1-K411R and RioK1-K411R/T410E in CRC cells promoted lung metastasis of CRC (Figure 9H). Compared to RioK1-WT, stable expression of RioK1-T410A in RKO cells caused a significant reduction of migration, invasion, and metastasis (Figure 9E, F, G, and H).

Additionally, all the *in vitro* and *in vivo* functional experiments in Figure 10 were repeated in MKN45 cells stably transfected with RioK1-T410A, RioK1-K411R/T410E, RioK1-T410A, and RioK1-WT, and showed similar results (data not shown). Together, these data confirm that the methylation and phosphorylation RioK1 oppositely regulates CRC tumorigenesis and metastasis.

### Clinical relevance of RioK1, SETD7, LSD1, FBXO6 and CK2 Expression in patients with CRC

To determine the clinical relevance of our findings, we performed immunohistochemical analysis of 104 primary CRC samples with validated antibodies recognizing RioK1, SETD7, LSD1, FBXO6, and CK2 proteins. Consistently, we observed an inverse expression pattern between RioK1 and SETD7 or FBXO6, whereas a positive expression pattern between RioK1 and LSD1 or CK2 was observed (Figure S5A and B). Kaplan–Meier analysis shows that high levels of LSD1 and CK2 in CRC samples correlated with poor overall survival in the patients, whereas low levels of SETD7 and FBXO6 in CRC samples correlated with poor overall survival (Figure S5C and D). Therefore, expression levels of these proteins were independent prognostic factors for overall survival and disease free survival. Together, our data highlight a critical role for RioK1 phosphorylation and methylation in the clinical aggressiveness of human CRC (Figure S5E).

## Discussion

In the present study, RioK1 appears to be important for CRC and GC progression because blockade of RioK1 decreases tumor growth and metastasis, whereas RioK1 overexpression causes the opposite effects. Furthermore, to our knowledge, we for the first time reveal that RioK1 is specifically monomethylated at K411 site by SETD7 *in vitro* and *in vivo*. This SETD7-mediated methylation recruits FBXO6 E3 ligase and significantly contributes to RioK1 instability. However, LSD1-mediated demethylation of RioK1 at K411 or CK2-mediated phosphorylation of RioK1 at T410 reverses RioK1 degradation by FBXO6. Importantly, functional experiments indicate that the RioK1 demethylation or phosphorylation signals might be a prerequisite for enhanced CRC and GC growth and metastasis. Clinically, PTMs of RioK1 are associated with CRC and GC patient prognosis. Together, these results provide a novel mechanism through which the crosstalk of PTMs regulates the function of RioK1, supporting the notion that RioK1 may serve as a valuable biomarker to monitor human CRC and GC development.

The most striking finding is that a methylation-phosphorylation switch dictates RioK1 stability and function in CRC and GC. Independent studies have validated the importance of RioK1 in cancer cell survival. A cell-based RNAi screen identified RioK1 required for Ras-mediated cell survival, although this study did not explored the functionality of RioK1 (Breitkreutz et al., 2010). Additionally, RioK1 upregulation has been shown to be positively associated with Akt activity in both glioblastoma specimens and cultured cell (Read et al., 2013). Consistent with these notions, in this manuscript, we demonstrate that RioK1 is upregulated in CRC and GC tumor cells relative to normal control cells, and RioK1 deficiency decreases tumor cell proliferation, migration, and lung metastasis. Therefore, RioK1 is a powerful modifier of tumorigenesis. Despite about one decade of study, very little was known about RioK1 regulation. It has been previously shown that Akt signaling regulates RIO kinase protein stability (Read et al., 2013), but the exact mechanism by which Akt regulates RioK1 levels remains undetermined. We now provide a novel perspective on the posttranslational regulation of RioK1 by showing that SETD7-mediated methylation of RioK1 decreases RioK1 protein stability, which can be reverse by LSD1. Moreover, we provide ample evidence that CK2-mediated phosphorylation of RioK1 at K410 reverses RioK1 degradation by FBXO6 and SETD7. Our data are the first to establish functional connections between the PTMs of RioK1 in CRC and GC. These results may have broad relevance to other cancers since RioK1 is strongly expressed in other tumor types (Faraji et al., 2014). Given that RioK1 creates a feedforward loop that promotes and maintains Akt activity (Read et al., 2013), we hypothesize that there might be a feedback loop between RioK1 and SETD7 or CK2, and that disruption of these loops are sufficient to trigger apoptosis or chemosensitivity in CRC and GC. This hypothesis awaits substantiation in future studies, which may lead to important new insights into the RioK1signaling network in both normal and cancer cells.

Compared with the plethora of information about histone methylation, our understanding of non-histone protein methylation is very limited. Although RioK1 has been shown to directly associates with the protein arginine methyltransferase 5 (PRMT5), RioK1 was not a substrate of PRMT5 (Guderian et al., 2011). RioK1 just acts as an adapter protein by recruiting the RNA-binding protein nucleolin to the PRMT5 complex for its symmetrical methylation (Guderian et al., 2011). In the present study, for the first time, we provide evidences that SETD7 and LSD1 methylates or demethylates RioK1 respectively. One of main mechanisms though which lysine methylation regulates protein function is effects on protein–protein interaction (Kim et al., 2016; Couture et al., 2006). Indeed, we observed that K411 methylation of RioK1 promotes the recruitment of FBXO6 to RioK1 and disrupts the association of CK2 and LSD1 with RioK1, subsequently leading to RioK1 degradation. Given that the level of RioK1 is closely related to diverse cancers (Read et al., 2013; Guderian et al., 2011), our data highlight the significance of understanding this important enzyme in health and disease.

Interplay with other PTMs is another main mechanism though which lysine methylation regulates protein function (Kim et al., 2016; Couture et al., 2006). Therefore, we also examined the phosphorylation and ubiquitnation of RioK1 by CK2 and FBXO6 respectively, which is the second important discovery in this study. Notably, CK2 was previously identified as an interaction partner of the Rio 1p, a homologs of the human RioK1 in yeast by MS analysis, and phosphorylation by CK2 leads to moderate activation of Rio1p in vivo and promotes cell proliferation (Estève et al., 2011), which is consistent with our data in human CRC and GC. Moreover, we now extend these findings by showing that CK2-mediated phosphorylation of the RioK1 promoted CRC and GC growth and metastasis by disrupting SETD7-mediated methylation of RioK1and recruitment of FBXO6 E3 ligase to RioK1. However, In that study, Six C-terminally located clustered serines were identified as the only CK2 sites present in Rio1p in yeast, and phosphorylation by CK2 renders the Rio 1p susceptible to proteolytic degradation at the G (1)/S transition, whereas the non-phosphorylated version is resistant (Estève et al., 2011), which stands in contrast with our current data. Notably, at first, the C-terminal domain of yeast Rio1p is least conserved in evolution, and especially lacks consensus SETD7 motif. Secondly, RioK1 of higher eukaryotes almost does not contain all six serine residues or CK2 motifs in the corresponding aligned sequences, but has highly conserved threonine residue and positively charged lysine residue next to it. This may provide an explanation for the most puzzling aspects of effect of CK2 on RioK1 in yeast and higher eukaryotes.

Recently, it has been demonstrated that LSD1 physically interacts with endogenous CK2 *in vivo* and is potentially phosphorylated by CK2 on three serine residues (S131, S137, S166) *in vitro* (34). And S137 facilitates RNF168-dependent ubiquitination and recruitment of 53BP1 to the DNA damage sites, which promotes cell proliferation and survival in response to DNA damage (Peng et al., 2015). Thus, it is possible that there exists a feedback loop between LSD1 and CK2 in CRC and GC. Additionally, it needs to be further clarified whether CK2-mediated phosphorylation of the RioK1 at Thr410 enhances the association of LSD1 with RioK1, or LSD1-mediated demethylation of RioK1 increases the binding of CK2 to RioK1 and phosphorylation.

Previous study suggested that the proteasome has a pivotal role in the regulation of RioK1 degradation (Read et al., 2013). However, no specific ubiquitin complex was previously connected to RioK1 ubiquitination. Although we can not formally rule out the contribution of other E3 ligase –mediated effects on the functions of RioK1, we were able to show here that FBXO6 links the activated RioK1 to the degradation machinery and FBXO6 activity affects the cancer-supportive role of RioK1 in CRC and GC. Importantly, we show that the ubiquitnation of RioK1 is controlled by molecular pathways (CK2 and LSD1and SETD7) frequently deregulated in CRC and GC (Peng et al., 2015; Lin et al., 2011; Akiyama et al., 2011). In the future, the identification of critical ubiquitination sites will provide new insights into the role and interaction of RioK1 in cancer.

Clinically, LSD1 and CK2 activation are associated with poor outcome in cancer (Peng et al., 2015; Lin et al., 2011). Interestingly, reduced SETD7 expression or inactivating mutations is significantly correlated with poor patient prognosis in multiple cancers (Akiyama et al., 2011). Recently, FBXO6 has been shown to affect the DNA damage, a process which promotes cell proliferation and survival in response to DNA damage (Zhang et al., 2011). Our data unify these observations and demonstrate that RioK1 accumulation due to altered SET79, LSD1, CK2, and FBXO6 expression provides an advantage in CRC and GC cells during disease progression. Indeed, we show a direct correlation between RioK1 and LSD1, CK2, FBXO6, or SETD7 protein expression to increased metastatic potential and decreased recurrence-free survival of CRC and GC patients. These data suggest that RioK1 could be a biomarker and a therapeutic target in these diseases. However, it remains to be seen whether altered function of these enzymes (SET79, LSD1, CK2, and FBXO6) in different cancers indeed converges on RioK1 stability.

In summary, we show that RioK1 is up-regulated in CRC and GC, which results in a significant reduced survival and a more malignant phenotype by enhancing proliferation and migration of CRC and GC cells. Most importantly, we show for the first time that methylation and phosphorylation control the stability of RioK1 and its role in CRC and GC progression. Therefore, targeting these PTMs of RioK1 could be a very promising novel treatment option for CRC and GC.

## Materials and Methods

Please see the supplementary materials and methods for detailed experimental procedures.

## Materials and Methods

### Cell culture and transfection

Colorectal cancer cell lines LOVO, RKO, M5, LS174T, HCT116, DLD1, SW480, SW620, HEK293T, MKN45 and SGC-7901, MEFs and a normal human intestinal epithelial cells (IECs) were obtained from the Cell Bank of the Chinese Academy of Sciences (Shanghai, China). All cells were cultured in Dulbecco’s Modified Eagles Medium (DMEM, Switzerland) supplemented with 10% fetal bovine serum (FBS) (Gibco, USA) and maintained in standard conditions (5% CO_2_ and 95% atmosphere, 37°C). Using a lentiviral system, stably transfected cells were created. Lentiviral vectors encoding the human RioK1 gene and its mutants or shRioK1, or empty vector (LV-control), or Non-specific shRNA were transfected into CRC or GC cells with a multiplicity of infection (MOI) of 40 to 50 in the presence of polybrene (5 μg/ml). At 48 hours after transfection, transfected cells were selected for 2 weeks with 2.5 μg/ml puromycin (Sigma). Pooled populations of knockdown cells and overexpression cells, which were obtained 2 weeks after drug selection without subcloning, were used in both in vitro and in vivo experiments.

### Quantitative PCR

Total RNA was extracted from tissues or cells using TRIzol reagent (Takara) according to the manufacturer’s instructions. RNA reverse transcription to cDNA was performed with a Reverse Transcription Kit (Takara). Quantitative real-time PCR (qRT-PCR) analyses used SYBR Green I (Takara) in triplicate. The results were normalized to the expression of GAPDH. The primer sequences used for qRTPCR are listed as follows: Human SETD7: 5′-AGTTCTCCAGGGCACGTATG-3′ (forward); 5′-TCTCCAGTCATCTCCCCATC5′-3′ (reversed) (SETD7); Human GAPDH: 5′-ACA GTC AGC CGC ATC TTC TT-3′ (forward) and 5′-GAC AAG CTT CCC GTT CTC AG-3′ (reversed); *Lsd1* forward: 5′-CGGCATCTACAAG AGGATAAAACC-3′, *Lsd1* reverse: 5′-CGCCAAGATCAGCTACATAGTTTC-3′. Indicated genes’ mRNA expressions were normalized to GAPDH level.

### Co-immunoprecipitation

Cells or CRC or normal tissues lysed in RIPA buffer (1 × PBS, 1% NP40, 0.1% SDS, 5 mM EDTA, 0.5% sodium deox ycholate, and 1 mM sodium orthovanadate) containing protease inhibitors followed by centrifugation at 14,000 rpm at 4 °C for 10 min. Total proteins (700 μg/sample) were pre-cleaned with 20 μl A/G beads (Santa Cruz) before immunoprecipitation with 10 μl control IgG (Bio-Rad), indicated primary antibodies overnight. After incubation with 20 μl A/G beads at 4°C for 12 h, the immunoprecipitates were washed with PBS containing 0.2% NP-40 for 5 times. The immunoprecipiated protein complexes were then released by boiling in 2×SDS-PAGE sample buffer for 5 min, and probed with indicated primary antibodies antibodies (Santa Cruz).

### cDNA constructs and RNA interference

The full-length open reading frames of RioK1 and SETD7 was amplified by PCR using the following primers: RioK1,5′-CGGACGTCGACATATGGACTACCGGCGGCTTCTCATG-3′ and 5′-CTGATGCGGCCGCCTATTTGCCTTTTTTCGTCTTGGC-3′ and confirmed by DNA sequencing. Target DNA fragments were then subcloned into pEGEX6P-1 (GE Healthcare) and pHA vector (, Invitrogen). Different truncated fragments of RioK1 were generated by PCR and subcloned into pEGEX6P-1 and pHA, respectively. The primer for cloning different truncations was as following, (aa 1–120, 5′-GTAGAATTCATGGACTACCGGCG-3′ and 5′-CATCTCGAGTCAATTAATTTTATTCTC-3′; aa 121–242, 5′-GATGCGGCCGCAATTTAGATAAGC-3′ and 5′-CATCTCGAGTCACATTTTCCTAGGGT-3′; and aa 1–242, 5′-CGGACGTCGACATATGGACTACCGGCGGCTTCTCATG-3′ and 5′-CGTGCGGCCGCTCACATTTCCTAGGG-3′), which was described as previously reported (1). Three RioK1 Human shRNA lentiviral constructs (OriGene, <u>TL320739</u>) were purchased from OriGene (USA). shRNA-resistant RioK1Δ was made by performing site –directed mutagenesis to introduce five silent mutations (CDS area 551 A > T, 554 T > A, 557 C > T, 566 T > C and 569 T > A) against RioK1 shRNA, primer sequences were listed as follow：Forward sequence:5'-CGG TTG ATA TTG ATT CCG CCA AAA CTC-3'and Reverse sequence: 5'-GAG TTT TTG TGA CGG AAT CAA TAT CAA-3'. The knocking effect of RioK1 was evaluated by western blotting using relevant specific antibodies. The siRNA oligonucleotide duplexes targeting CK2α (sequence: GAUGACUACCAG CUGGUUCdTdT), CK2α’ (sequence: CAGUCUGAGGAGCCGCGAGdTdT), LSD1 3′ UTR (sequence: GCUCUUCUAGCAAUACUAGdTdT). Oligonucleotides the shRNA sequences were as following. For silencing of SETD7: GCAAACTGGCTACCCTTATGT (shSETD7–1); and GGGAGTTTACACTTACGAAGA (shSETD7–2). The shcontrol and sh-LSD1 (shLSD1–1: CACCGCCTGTTTCTGC CATGTAAGGCGAACCTTACATGGCAGAAACAGGC, shLSD1–2: CACCGCCAT GTAAGGAAGGCTCTTCCGAAGAAGAGCCTTCCTTACATGGC) Specific siRNAs and scrambles were purchased from IBA Nucleic Acids Synthesis, Germany) and transfected with Oligofectamine^TM^ (Invitrogen) according to the manufacturer’s protocol. The knocking effect of RioK1 was evaluated by western blotting using relevant specific antibodies. SETD7-H297A and RioK1-K411R and K413R mutants were introduced using the QuikChange site-directed mutagenesis kit (Stratagene) according to the manufacturer’s instructions. All of the constructs were confirmed by DNA sequencing. The pcDNA-myc/his-SETD7 was created by subcloning SETD7 into pcDNA3.1/myc-His A (Invitrogen). SETD7 siRNA (Ambion, siRNA sequence: 5′‐ AGAUAACAUUCGUCAUGGA‐ 3′) or non‐ targeting siRNA (Dharmacon). The plasmid for V5-SETD7 was generated by PCR cloning. Flag-tagged SETD7-WT or -H297A FL and fragment cDNAs were amplified from plasmids by PCR and separately subcloned into pGEX-4T3, EGFP, and pET28 vectors. Human cDNA clones encoding LSD1, CK2α, CK2α’ and CK2β were subcloned into pcDNA3.0 backbone as well as with three copies of HA or FLAG tag at the N-terminus or one copy of MYC tag at the C-terminus. The expression construct of HA-Ub was a gift from Dr. Bing Wang (Rutgers University).

### *In vivo* ubiquitination assay

Indicated CRC or HEK293T cells were transfected with combinations of plasmids including His-ubiquitin. After 48 h, they were incubated with MG132 (10 μg/ ml) for 6 h, lysed in buffer A (6 M guanidium-HCl, 5 mM imidazole, 0.1 M Na_2_HPO_4_/NaH_2_PO_4_, 0.01 M Tris-HCl pH 8.0, and 10 mM β-mercaptoethanol), and at room temperature treated with Ni^2+^-NTA beads (Qiagen) for 4 h. The beads were washed sequentially with buffer A, buffer B (0.01 M Tris-Cl pH 8.0, 10 mM β-mercaptoethanol, 8 M urea, and 0.1 M Na_2_PO_4_/NaH_2_PO_4_), and buffer C (8 M urea, 0.1 M Na_2_PO_4_/NaH_2_PO_4_, 0.01 M Tris-Cl pH 6.3 and 10 mM β-mercaptoethanol). Bound proteins were eluted with buffer D (30% glycerol, 0.72 M β-mercaptoethanol, 200 mM imidazole, 0.15 M Tris-Cl pH 6.7, and 5% SDS), finally were analyzed with Western blot.

### Cell proliferation

The cell proliferation assay was performed with the Cell Counting Kit-8 (CCK-8, Sigma) or MTT assays according to the manufacturer’s instructions.

### *In vitro* methylation and demethylation assays

GST, GST-RioK1, His-SETD7, or His-SET7-H297A were expressed in BL21/pLysS cells and purified using nickel-nitrilotriacetic acid, or glutathione-Sepharose 4B (GE Healthcare), or in the form of GST fusion SETD7 was purified from *E.coli*. We performed *in vitro* methylation assays using 30 μl reaction volume as previously reported (1). Birefely, add 2 μg of recombinant SET7 or SET7-H297A as well as with 2 μg of recombinant RioK1 proteins in the presence of 0.1 mm *S*-adenosylmethionine (Sigma, A2408) in the reaction buffer (4 mm DTT, 5 mm MgCl_2_, 50 mm Tris-Cl, pH 8.5) at 30 °C for 1 h. The reaction was stopped by boiling in 2× SDS-sample buffer, analyzed by Coomassie staining and radiography of the dried gel, or analyzed by Western blotting.

### Migration and invasion assays

6.5-mm diameter Boyden chambers with pore size of 8.0 μm (Corning) was used for migration assays. Briefly, the stable cell lines (3.0 × 10^5^ cells per well) were resuspended in the migration medium without FBS, and placed in the upper compartment of transwell chambers. The lower compartment was filled with 500 μl medium containing 10% FBS as a chemoattractant. Cells were fixed in 4% formaldehyde, and stained with 0.1% crystal violet after 24 hours. Ten random fields were counted under a light microscope (Carl Zeiss, Germany). However, cell invasion (3.0 × 10^5^ cells per well) was evaluated in 24-well matrigel-coated invasion chambers after 48 hours.

### GST pulldown assay

GST or GST-fusion proteins were expressed in *Escherichia coli,* and induced by 0.5 mm IPTG (isopropyl-β-d-thiogalactopyranoside) at 37°C for 3 h, and purified by glutathione-Sepharose 4B beads (GE Healthcare, USA), and then washed with TEN buffer [0.1 mM EDTA, 100 mM NaCl, and 20 mM Tris-HCl (pH 7.4)]. Recombinant His-tagged proteins were purified from *E. coli* by Ni (ii)-Sepharose affinity (GE Healthcare). Purified GST-tagged RioK1, and its truncations, and 2 μg of His-SETD7, and GST (around 10 μg) were cross-linked to glutathione-Sepharose by dimethyl pimelimidate dihydrochloride, and incubated with indicated cell extract in reaction buffer (1 mm EDTA; 0.1% (v/v) Nonidet P40; 20 mm HEPES, pH 8.0; 150 mm KCl; 10% (v/v) glycerol) for 2 h at 4 °C. The matrix was washed and bound proteins were eluted by boiling in 2× SDS sample buffer, and then analyzed by Coomassie staining and Western blot.

### Purification of RioK1-binding proteins

RioK1-binding proteins were purified from extracts of HEK293T cells expressing Flag-tagged RioK1. Mock purification from HEK293T cells expressing an empty vector was performed as a negative control. The procedure has been described previously (2). In brief, The RioK1-binding proteins were precipitated using Flag M2 agarose beads (Sigma, 100 μl of 50% slurry) from ~100 mg of cell extracts. After overnight incubation at 4 °C, the beads were washed three times with a BC150 buffer (1 mM EDTA, 0.05% Nonidet P40, 1 mM dithiothreitol, 0.2 mM PMSF, 20 mM Tris-HCl pH 7.9, 15% glycerol, and 150 mM KCl), and twice with a BC300 buffer, and three times with a TBS buffer. Finally, the bound proteins were eluted, and resolved by SDS–PAGE for LC-MS/MS analysis.

### Mass spectrometry

Mass spectrometry was performed as described previously (3).

### *In vitro* ubiquitination

In vitro ubiquitination assay was performed according to previous report (4). In Summary, HA-RioK1 was expressed in 293T cells, pulled down with anti-HA antibodies, and the HA-RioK1-bound beads were used as the ubiquitination acceptor substrate. Extracts from indicated cells expressing Myc-FBXO6 were immunoprecipitated using protein A-agarose beads coupled with anti-Myc, and treated with hypotonic buffer (2 mM DTT, 0.25 mM EDTA, 20 mM Tris-HCl, pH 7.2, and protease inhibitors), Then we performed *in vitro* ubiquitination assay in one reaction buffer (5 mM MgCl2, 0.5 mM DTT, 50 mM Tris-HCl, pH 7.5, 2 mM NaF, 10 nM okadaic acid) with 30 μM MG132, 1 μg/ml recombinant Flag-Ub, and 10 μl concentrated indicated cell extracts.

### Western blotting and antibodies

Cells were harvested and lysed on ice for 30 min in 1× RIPA as described above. Cell lysates were analyzed by SDS-PAGE and transferred to PVDF memebrane. Membranes were washed and incubated with indicated antibodies at 4°C overnight respectively. Antibody against RioK1 was raised by injection of recombinant RioK1 comprising aa 1–242, and affinity-purified on columns with the respective covalently linked antigen. Then membranes were washed and incubated with horseradish peroxidase-conjugated secondary antibodies (BIO-RAD) according to the manufacturer’s instructions. The protein of interest was visualized using ECL Western blotting substrate (Pierce). Or samples were resolved by SDS-PAGE and further analyzed by silver staining and Coomassie staining. Other antibodies are: Anti-HA (F-7, Santa Cruz Biotechnology), anti-FLAG (M2, Sigma-Aldrich), ubiquitin from Santa Cruz, and V5 from Invitrogen. The homemade rabbit anti-SETD7 polyclonal antibody was raised against recombinant GST-SETD7 protein. The mouse anti-SETD7 monoclonal antibody was purchased from Millipore and Cell Signaling. Rabbit polyclonal antibodies used in this study including anti-HA (A190–208A), anti-MYC (A190–205A), anti-LSD1 (A300–215A), (A300–272A), anti-CK2α (A300–198A), anti-CK2α’ (A300–199A), anti-CK2β (A301–984A), anti-GAPDH (A300–643A) were purchased from Bethyl. RioK1-K119me, p-T410 and 400-414 antibodies were generated by immunizing rabbits with K411-monomethylated, T410-phosphorylated and unmodified 400-414 peptides conjugated with keyhole limpet hemacyanin (KLH), respectively. Briefly, rabbits were immunized by keyhole limpet hemocyanin–conjugated peptides. The sera were tested and collected after 4 more antigen boosters. To eliminate nonspecific recognition of nonmethylated antigen, the sera were precleaned by incubating with nonmethylated peptide and examining with affinity chromatography. The specific polyclonal antibodies were finally purified by methylated peptide affinity chromatography *via* CNBr-activated Sepharose 4 Fast Flow (17-0981-01; GE Healthcare). Anti-FBXO6 antibodies were generated against KLH-coupled synthetic peptide corresponding to amino acids 262 to 284 of human FBXO6, which lie very close to the C terminus. Antiserum from peptide-injected rabbits was affinity purified using the AminoLink Plus Immobilization Kit (Pierce).

### *In vitro* protein phosphorylation assays

*In vitro* kinase assays were performed as described before (5) except that the CK2 kinase buffer contained 1 mM DTT, 1 mM Na_3_VO_4_, 10 μM cold ATP, 50 mM HEPES (pH 7.4), 10 mM MgCl2, and 5 μCi [γ-^32^P]-ATP. The GST pulldown after kinase assay was performed by adding 500 μl of NETN buffer into the reaction after the *in vitro* kinase assay was completed. In all, 25 μg of RioK1 peptides were used in similar reactions prior to direct spotting onto nitrocellulose for immunoblot.

### Immunofluorescence

For immunofluorescent staining, indicated cells plated on poly-llysine-coated glass coverslips in DMEM and 10% FCS were washed once in PBS, then were fixed with 4% formaldehyde, permeabilized with 0.2% Triton X-100 and blocked in 1% BSA in PBS. cells were incubated for 1 h with indicated primary antibodies at 4°C overnight. Following staining with rhodamine-conjugated secondary anti-rabbit antibodies, cells were analyzed with a 63× oil immersion lens on a Zeiss Axiovert 200 m microscope. Indicated cells transfected with indicated plasmids were directly analyzed after 48h without fixation. The fluorescent staining was recorded using an inverted fluorescence microscope (Leica, Germany).

### Patients specimens and tissue microarray

From January 2008 to May 2014, a total of 104 pairs of CRC and 20 pairs of GC and non-tumor samples were collected from patients who had CRC or GC resection performed at Third Affiliated Hospital of Harbin Medical University. The samples were used for subsequent RNA extraction or immunohistochemistry (IHC) stained for RioK1, FBXO6, CK2, LSD1, SETD7. All human materials were obtained with informed consent and approved by the ethics committee of Hospital of Harbin Medical University. The tissue specimens were frozen in liquid nitrogen and stored at −80 °C. All tissues were confirmed as adenocarcinoma. CRC tissue microarray (TMA) was purchased form US Biomax (BC051110b) and IHC stained for RioK1. The slides were scored for the percentage of positive epithelial cells as well as intensity and protein staining was subsequently quantified using the a scoring system based on the Allred score for immunohistochemistry (6,7). Based on the RioK1protein expression as defined by Allred score <6 (low) and >6 (high), patients were stratified into two groups: RioK1 low and high expression groups.

### In vivo tumorigenic and metastasis assay

All animal experiments were performed in accordance with NIH guidelines for the use of experimental animals. Male nonobese/severe combined immunodeficiency (NOD/SCID) mice between 4 and 6 weeks of age, obtained from the Experimental Animal Center of Shanghai Institute for Biological Sciences (SIBS). All animal work was conducted according to national guidelines, and all animal experiments were approved by the ethical committee of the Harbin Medical University. HCT116-pLV-shRioK1 and HCT116-PLV-shNC and other indicated cells (1 × 10^7^ cells per mouse) were injected subcutaneously into the left or right flank of NOD/SCID mice (n = 5 per group). After 8 weeks, the remaining mice were sacrificed, and lungs were isolated for examination of the number of metastatic tumors. Tumor volume (V) was estimated from the length, width (w), and height (h) of the tumor. We analyzed primary tumor growth by the formula (length × width^2^)/2. For the metastasis model, 2 × 10^6^ cells were injected into the tail vein of NOD/SCID mice (n = 5 per group); eight weeks later, the mice were sacrificed, and their lungs were removed and formalin fixed for haematoxylin and eosin (HE) staining.

### Clonogenic assay

Clonogenic cell survival assay was performed to evaluate CRC and GC cells proliferation potential. In a 10cm dish, a single cell suspension (250 cells) was added to each dish. Two weeks after plating, single colonies were fixed and stained by haematoxylin. Then colonies were counted and measured using the Nikon Nis Elements^®^ for windows software. Each assay was performed three times in triplicate.

### Statistical analysis

All statistical analyses were performed using the SPSS 16.0 statistical software package (Abbott Laboratories, USA). Quantitative values of all experiments are expressed as the mean±SD. The significance of correlation between the expression of indicated proteins and histopathological factors was determined using Pearson χ^2^ test. Survival curves were plotted by Kaplan–Meier method and compared by log-rank test. In vitro cell growth assay was tested using factorial design ANOVA. Comparisons between groups were performed with a 2-tailed paired Student’s *t*-test. The correlation between the expression levels of two proteins was determined using Spearman’s correlation analysis. In all samples, *P* < 0.05 was considered to be statistically significant. Statistical significance was concluded at \**P* < 0.05, \*\**P* < 0.01, \*\*\**P* < 0.001; # represents no statistical significance.

**Supplementary Figure Legend 1.**
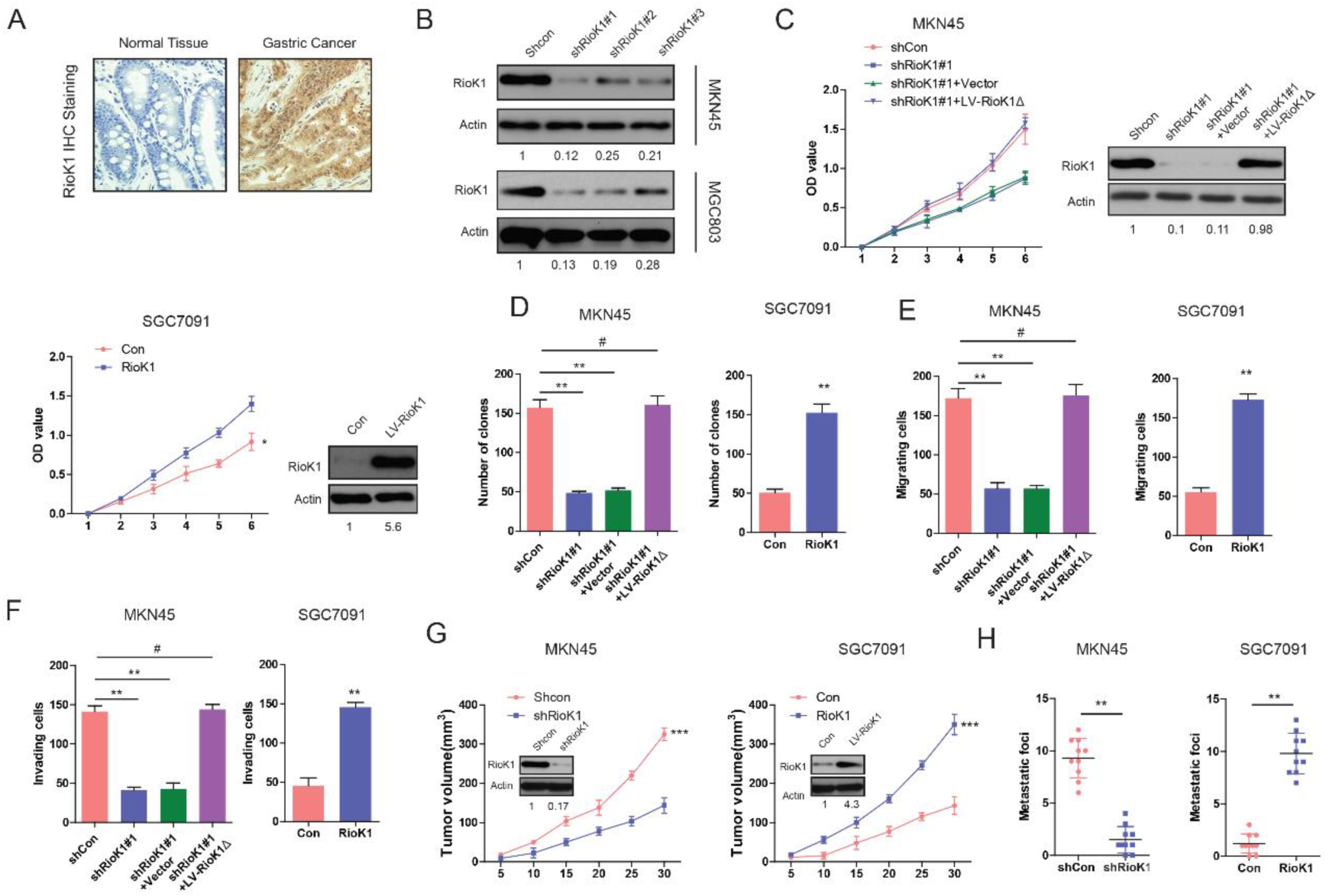
RioK1 promotes the proliferation, invasion and metastasis of GC *in vitro* and *in vivo.* (A) Immunohistochemical analysis of RioK1 on a group of patients with GC (n=20) and healthy adjacent tissue (n=20. (B) Western blot for RioK1 in MKN45 cells infected with shRioK1, or control shRNA lentiviral vector, as well as in SGC7091 infected with the RioK1-expressing or empty control vector (n = 4). And the intensity of the western blot bands was quantified using NIH ImageJ software.(C). Western blot for RioK1 in the indicated MKN45 cells transfected with shRioK1#1 or the shRNA-resistant expression construct, RioK1Δ. RioK1 expression was recovered in the MKN45-shRioK1 cells transfected with RioK1Δ. And the intensity of the western blot bands was quantified using NIH ImageJ software. RioK1Δ almost restored the proliferative ability of the cells. The effect of RioK1 on proliferation of MKN45 and SGC7091 cells was determined by CCK-8 and (D) colony formation assays. (E and F) The effect of RioK1 on migration and invasion of MKN45 cells was determined by transwell assays. (G) Growth curve of subcutaneous injection of indicated plasmids in NOD/SCID mice (n=6 per group). The mice were sacrificed and the tumors were then removed, weighed and compared. And knockdown or overexpression of RioK1 in a xenograft tumor model was analyzed with western blotting, and the intensity of the western blot bands was quantified using NIH ImageJ software. (H) Effects of RioK1 knockdown or over-expression on tumor metastasis of indicated cells in NOD/SCID mice (*n* = 10 per group): the number of metastatic nodules in the lung. The results are presented as are means ± SD (*n* = 3 for each panel). Statistical significance was concluded at \**P* < 0.05, \*\**P* < 0.01, \*\*\**P* < 0.001; # represents no statistical significance.

**Supplementary Figure Legend 2.**
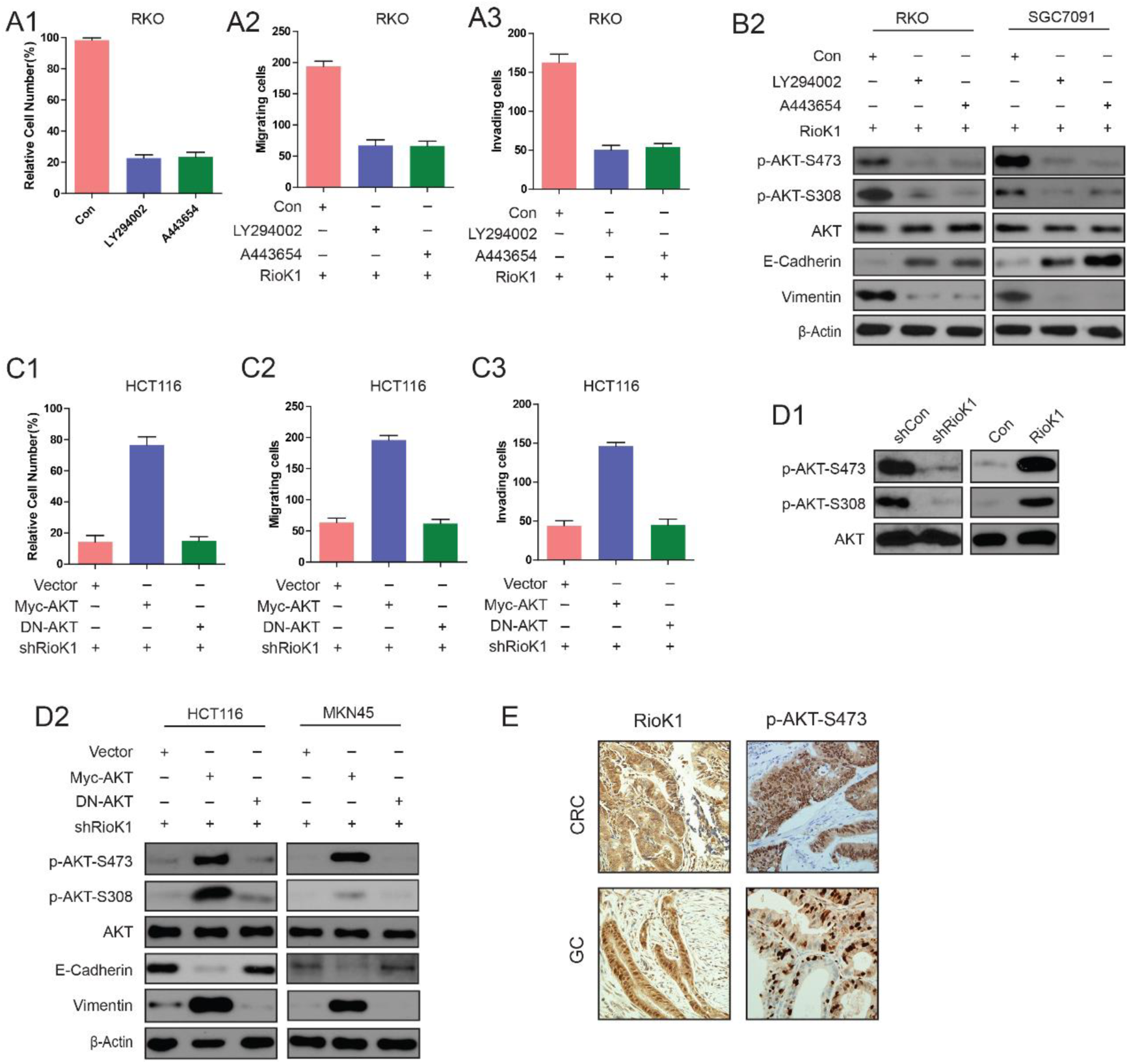
PI3K/AKT pathway is required for RioK1-mediated CRC and GC cell proliferation and migration. (A) CRC and GC cells were transfected with RioK1 plasmid, and treated with LY29004 (30μM) or A443654 (1 μM). Then cell proliferation (A1), migration (A2) and invasion (A3) were determined. (B) The whole-cell lysates in (A) were analyzed by Western blot with indicated antibodies. (C) The RioK1-knockdown-HCT116 and MKN45 cells were transfected with indicated plasmids. The cell proliferation (C1), migration (C2) and invasiveness (C3) was tested. (D1 and D2) The cells in (C) were harvested at 48 hr for western blot analysis with indicated antibodies. The data represent the mean ± SEM from three independent experiments. (E) Immunohistochemistry analysis for RioK1 and phospho-AKTSer473 in human CRC and GC tissues.

**Supplementary Figure Legend 3.**
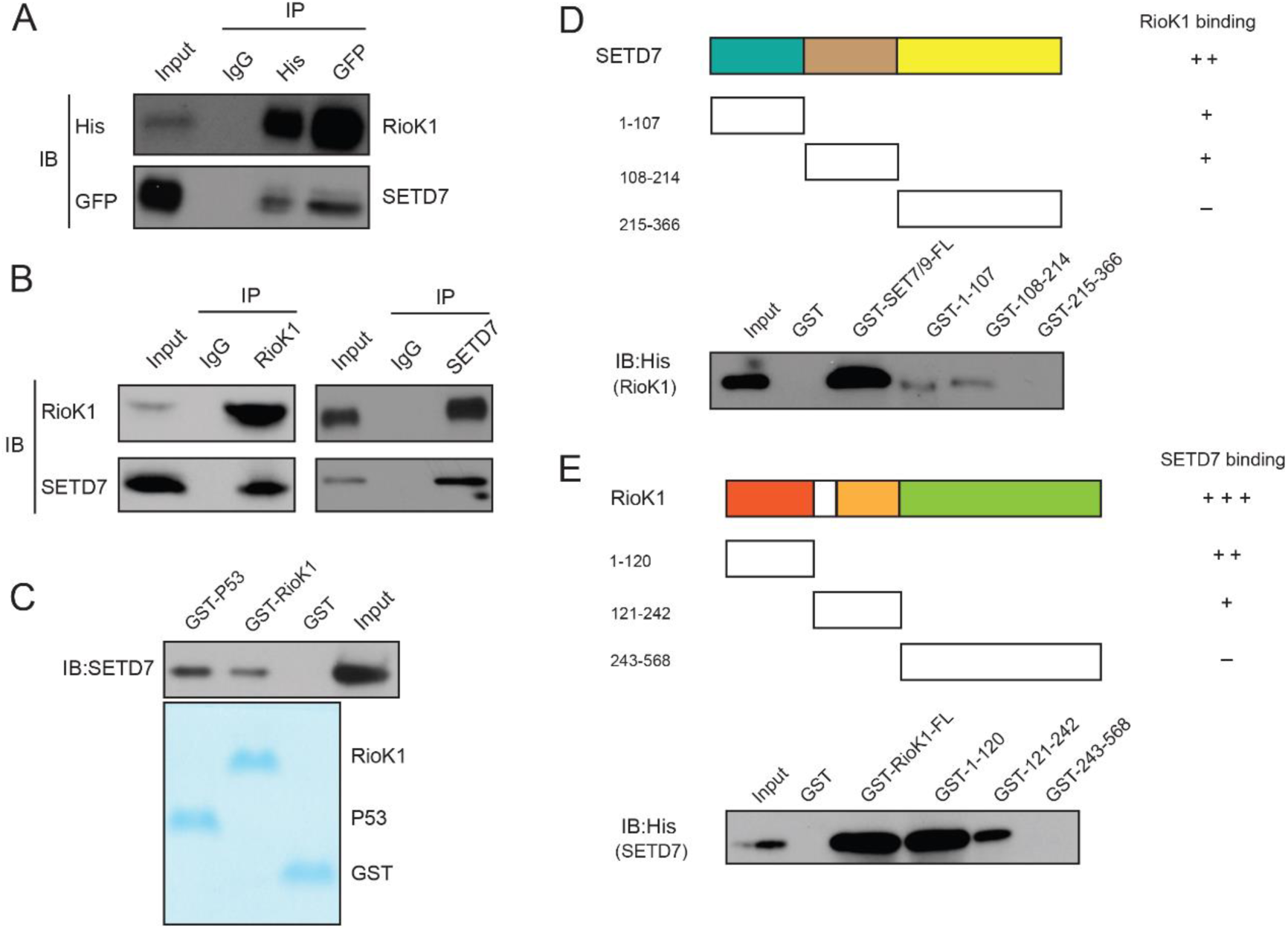
SET7/9 directly interacts with RioK1 *in vitro* and *in vivo*. (A) Whole-cell lysates of RKO cells transfected with GFP-SET7/9 and His-RioK1 were precipitated with an anti-GFP or anti-His antibody, and the interactive components were analyzed by Western blot. (B) RKO cells were extracted and immunoprecipitated with an anti–RioK1 (left) or anti-SET7/9 (right) antibody. IP with rabbit IgG was used as the negative control. Western blot analysis was performed with the antibodies indicated. (C) GST (negative control), GST-RioK1, or GST-p53 (positive control) was incubated with whole-cell lysates of HeLa cells, and Western blot analysis was performed to detect the interaction between SET7/9 and RioK1. (D) GST-SET7/9 FL or fragments were incubated with His-RioK1, and Western blotting was performed to detect the interaction with an anti-His antibody. (E) The reciprocal pull-down assay between GST-RioK1 FL or fragments and His-SET7/9.

**Supplementary Figure Legend 4.**
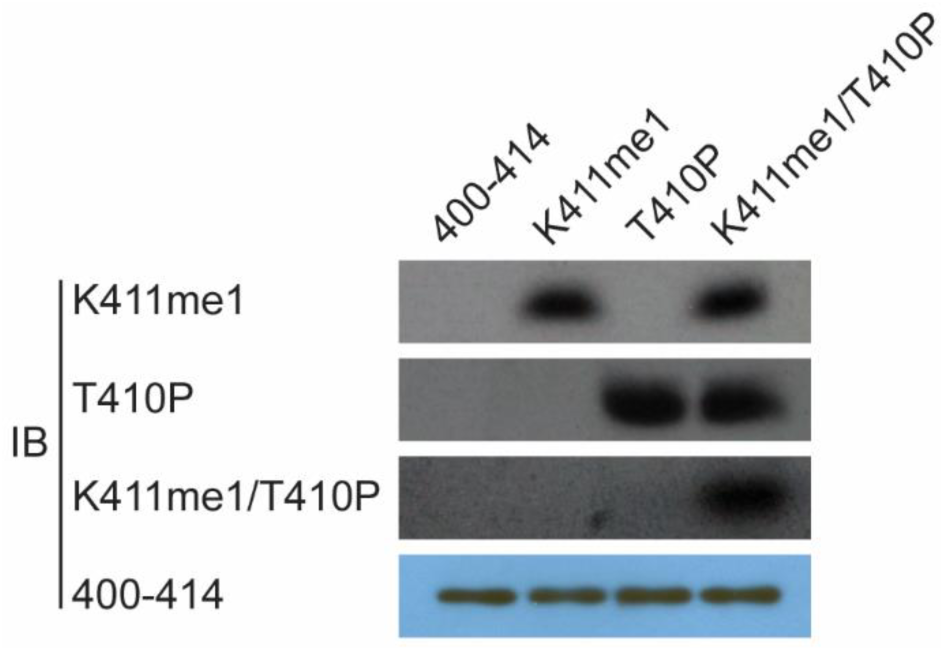
The validation of the antibodies’ specificity. To easily find whether methylation of RioK1 occurred in cells, we prepared a modification-specific antibody, anti-RioK1-K411me1, which specifically recognized monomethylated RioK1-K411 but not the unmethylated peptide. This antibody also recognizes a pT410.K411me1 doubly modified peptide as efficiently as the K411me1 peptide. Conversely, the anti-pT410 antibody recognizes a pT410.K411me1 peptide as efficiently as the pT410 peptide. These specific antibodies are characterized. RioK1 peptide substrates: 400-414 SKAMEIASQRTKEER, K411me SKAMEIASQRTKmeEER, T410p SKAMEIASQRTpKEER, K411me/T410p SKAMEIASQRTpKmeEER, p: phosphorylation; me: methylation.

**Supplementary Figure Legend 5.**
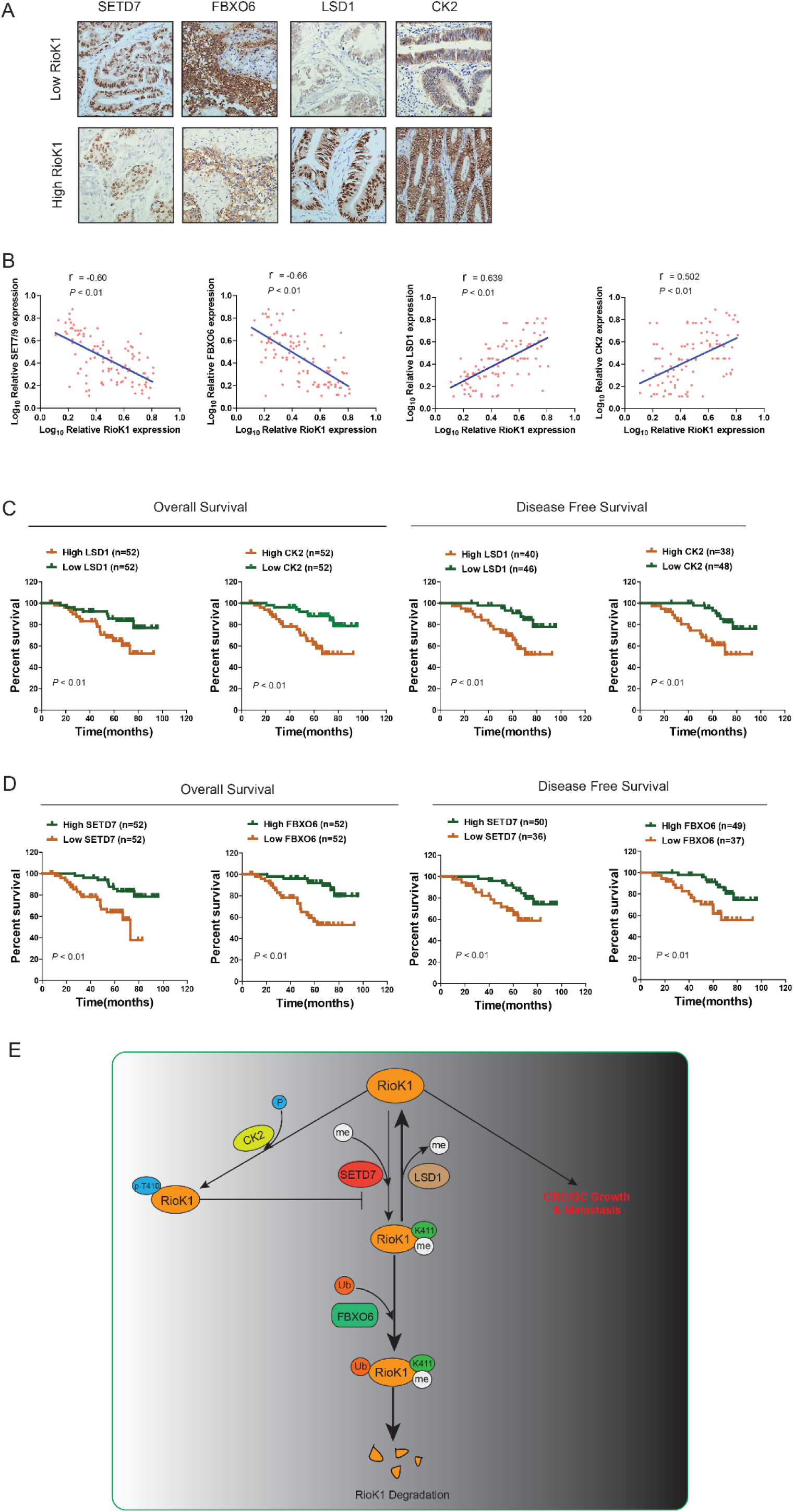
Clinical relevance of RioK1, SET7/9, LSD1, FBXO6 and CK2 Expression in patients with CRC. (A) Immunohistochemical staining of 104 human CRC for LSD1, CK2, SET7/9, and FBXO6 was performed. Representative photos of stains are shown in the groups with high (staining score, 5–8.0) and low (staining score, 0–4.0) expression of RioK1. (B) Correlation between RioK1 expression and LSD1, CK2, SET7/9, and FBXO6 expression in 104 clinical samples, respectively. (C and D) Kaplan–Meier plots of the overall survival and disease free survival in the patients (*n* = 104) with CRC in the groups with high (staining score, 5–8.0) and low (staining score, 0–4.0) expression of LSD1, and CK2 and SET7/9 and FBXO6. (E) A proposed model illustrating that the RioK1 methyl-phospho switch by SET7/9- CK2-LSD1 axis dictates the stability of RioK1 and its role in CRC and GC growth and metastasis

---

**Supplementary Table 1.** Univariate analysis and multivariate analysis between RioK1 expresssion and Clinicopathologic Features of CRC Patients (n=104)

| Variables | Univariate analysis | Multivariate analysis |  |
| --- | --- | --- | --- |
|  | *P* | HR (95% CI) | *P* |
| **Age** |  |  |  |
| <65 | 0.129 |  |  |
| >65 |  |  |  |
| **Gender** |  |  |  |
| Male | 0.510 |  |  |
| Female |  |  |  |
| **Depth** |  |  |  |
| T1/T2/T3 | **0.030** | 2.449 | 0.054 |
| T4 |  | (0.986-4.983) |  |
| **Lymph node metastasis** |  |  |  |
| Negative | **0.001** | 4.876 | **0.001** |
| Positive |  | (2.386-8.820) |  |
| **Venous invasion** |  |  |  |
| Negative | **0.034** | 2.262 | 0.146 |
| Positive |  | (0.753-4.796) |  |
| **CEA** |  |  |  |
| <5 | **0.008** | 3.177 | **0.019** |
| >5 |  | (1.212-6.329) |  |
| **Lymphatic invasion** |  |  |  |
| Negative | **0.022** | 1.548 | 0.458 |
| Positive |  | (0.488-4.909) |  |
| **RIOK1** |  |  |  |
| Low expression | **0.006** | 4.521 | **0.003** |
| High expression |  | (1.648-8.402) |  |

**Supplementary Table 2.** Identification of RioK1-interacting proteins via MS

Identification of RioK1-interacting proteins via MS
| Swiss-Prot accession no. | Gene Symbol | Protein name | No. of unique peptides matched | % Sequence identified by MS/MS data | Protein mascot score |
| --- | --- | --- | --- | --- | --- |
| Q969H0 | FBXW7 | F-box/WD repeat-containing protein 7 | 9 | 29 | 192 |
| Q9NRD1 | FBXO6 | F-box only protein 6 | 6 | 23 | 179 |
| Q13547 | HDAC1 | Histone deacetylase 1 | 7 | 18 | 168 |
| P61978 | HNRNPK | Heterogeneous nuclear protein K | 5 | 25 | 154 |
| O14744 | PRMT5 | Protein arginine N-methyltransferase 5 | 6 | 21 | 143 |
| P31946 | YWHAB | 14-3-3 protein beta/alpha | 6 | 18 | 139 |
| Q13573 | SNW1 | SNW domain-containing protein 1 | 7 | 19 | 127 |
| P61313 | RPL15 | 60S ribosomal protein L15 | 5 | 17 | 121 |
| Q9ULK4 | MED23 | Mediator of RNA polymerase II transcription subunit 23 | 4 | 20 | 107 |
| P11441 | UBL4A | Ubiquitin-like protein 4A | 4 | 15 | 95 |
| P00533 | EGFR | Epidermal growth factor receptor | 3 | 17 | 87 |
| Q15717 | ELAVL1 | ELAV-like protein 1 | 4 | 18 | 76 |
| O60341 | LSD1 | Lysine-specific histone demethylase 1A | 5 | 20 | 69 |
| P61956 | SUMO2 | Small ubiquitin-related modifier 2 | 3 | 16 | 51 |

## Notes

**Grant support:** This study was supported by National Natural Science Foundation of China (Grant No.81602149, No. 81172283, No.81141068, No.31571241), Natural Science Foundation of Fujian Province (Grant No. 2016J01619, No. 2012D037, No. 2015J01530), Training Program for Young Talents of Fujian Health System (Grant No. 2016-ZQN-85), Clinical Research Foundation of WU JIEPING Medical Foundation (No. 320.6750.13105).

**Conflict of interest:** The authors declare no conflict interest.

